# Label-free and reference region-free X-ray cross-β index for quantifying protein aggregates of neurodegenerative diseases

**DOI:** 10.1101/2024.01.31.578167

**Authors:** Karthika Suresh, Eshan Dahal, Aldo Badano

**Affiliations:** Division of Imaging, Diagnostics, and Software Reliability, Office of Science and Engineering Laboratories, Center for Devices and Radiological Health, Food and Drug Administration, Silver Spring, MD 20993, United States of America

**Keywords:** Neurodegenerative diseases, neuropathology, protein aggregate quantification, regional aggregate burden, X-ray scattering, material decomposition

## Abstract

**Background and objectives:** Recent advancements in therapies targeting various protein aggregates, ranging from oligomers to fibrils, in neurodegenerative diseases exhibit considerable promise. This underscores the imperative for robust quantitative methods capable of accurately detecting and quantifying these biomarker aggregates across different structural states, even when present in sparse quantities during the early stages of the disease continuum. In response to this exigency, we propose and assess an X-ray-based quantitative metric designed for the global and region-specific detection and quantification of oligomers and fibrils within tissues. This methodology proves applicable to a broad spectrum of neurodegenerative diseases, including Alzheimer’s and Parkinson’s. Notably, unlike positron emission tomography (PET)-based biomarker quantification methods, our approach obviates the need for a contrast agent or a reference region.

**Methods:** We assessed the proposed metric, termed X-ray cross-β aggregate index (XβAI), in a sheep brain model and brain tissue phantoms, incorporating synthetic oligomers and fibrils characterized against amyloid β-42 and α-synuclein aggregates. Detection of these biomarkers utilized laboratory-based monochromatic, and polychromatic X-ray sources, specifically targeting the cross-β substructure of protein aggregates. We employed a peak-location, knowledge-based material decomposition approach to extract target signals from the complex X-ray scattering spectrum originating from a mixture of tissue, water, and aggregate signals.

**Results:** Clinically relevant quantities of oligomers and fibrils were detected in tissues from different brain regions using the laboratory-based X-ray scattering method, without the need for a contrast agent. The signals from protein aggregates were successfully recovered from composite X-ray scattering spectra through material decomposition, eliminating the need for a reference region. The area under the peak of the decomposed inter-β-strand signal correlated well with aggregate burden in synthetically diseased brain tissues. The X-ray cross-β aggregate index (XβAI) accurately quantified aggregate burden in heterogeneous tissues across various brain regions and effectively tracked the deposition increments in specific tissue regions.

**Conclusion:** Our study introduces a novel metric, for both regional and global quantification of protein aggregates linked to various protein misfolding diseases, including synucleinopathies.

## 1. INTRODUCTION

In the realm of neurodegenerative diseases (NDDs), the standard clinical practice in diagnostic imaging is predominantly qualitative in nature. The U.S. Food and Drug Administration (FDA) approved diagnostic imaging agents that bind to specific pharmacologic target are mainly for qualitative analysis based on visual read (Table 1). For instance, the FDA-approved DaTscan (Ioflupane I 123 Injection) is for the striatal dopamine transporter visualization using single photon emission computed tomography (SPECT) brain imaging to assist in the evaluation of adult patients with suspected Parkinsonian syndromes (PS) [1]. To evaluate the PS suspected patients, another FDA-approved Fluorodopa F18 tracer helps to detect the damaged or lost dopaminergic nerve cells [2]. Similarly, the amyloid positron emission tomography (PET) radiotracers and [18F] Flortaucipir are for the visual rating of amyloid neuritic plaques and tau neurofibrillary tangles (NFTs) respectively for adult patients with cognitive impairment who are being evaluated for Alzheimer’s disease (AD) (Table 1) [3–5]. However, recent studies suggest that biomarker quantification in conjunction with visual reads can improve the diagnostic confidence for NDDs [6].

**Table 1:** The list of the FDA approved or select list of breakthrough device designation granted medical imaging agents or fluid biomarker assays for NDDs with a focus on AD, Parkinson’s disease (PD) and tauopathies.

| Pharmacologic target | Radiotracer/ligand for imaging | CSF<sup>‡</sup> | Plasma |
| --- | --- | --- | --- |
| **Amyloid-β** | [18F] Florbetapir (Amyvid™)<br>[18F] Flutemetamol (Vizamyl™)<br>[18F] Florbetaben (Neuroceq™) | Lumipulse G β-Amyloid Ratio (Aβ-42)/(Aβ-40)<br>Elecsys β-Amyloid (1-42) CSF II for Aβ42<sup>2</sup><br>Elecsys β-Amyloid (1-42) CSF II Elecsys Total-Tau CSF<sup>3</sup> | PrecivityAD™,<sup>4</sup> |
| **Tau** | [18F] Flortaucipir (Tauvid™,<sup>1</sup>) | Elecsys Phospho-Tau (181P) CSF for pTau181<sup>2</sup><br>Elecsys Total-Tau CSF<sup>3</sup> | Simoa ptau-181 Advantage V2 Kit<sup>4</sup><br>Elecsys® Amyloid Plasma Panel (EAPP)<sup>4,5</sup> |
| **α-synuclein** |  | SYNTap®,<sup>4</sup> |  |
| **Dopamine transporter imaging** | DaTscan™ Ioflupane I 123<br>Fluorodopa F18 |  |  |

<sup>1</sup>second generation tracer- tau pathology in AD. <sup>2</sup>common name is Elecsys Phospho-Tau (181P) Elecsys β-Amyloid (1-42) CSF II ratio, used together to estimate the ratio of p-tau181/Aβ42. <sup>3</sup>Elecsys Total-Tau CSF is used for the quantitative measurement of tTau in human CSF with the intent that the result only be used to calculate the ratio of Elecsys Total-Tau CSF to Elecsys β-Amyloid (1-42) CSF II. <sup>4</sup>not FDA approved but FDA granted breakthrough device designation. The FDA breakthrough devices Program is designed to speed up the development, assessment, and review of medical devices that provide more effective treatment or diagnosis of life-threatening or irreversibly debilitating diseases or conditions. <sup>5</sup>combines the result of the phosphorylated Tau (pTau) 181 protein assay and apolipoprotein (APOE) E4 assay in human plasma. <sup>‡</sup>cerebrospinal fluid.

Quantitative methods enable a continuous measurement of target levels, surpassing the dichotomous assessment of presence or absence provided by visual reads. Recent studies have found that the quantification could supplement visual inspection of target imaging in amyloid PET, especially for accurate early detection of biomarkers, more difficult to interpret cases like the equivocal or grey zone regions (PET centiloid scales, CL= 15-24) and for less experienced readers [6]. Additionally, the quantification enables the measurement of disease progression and the stratification of patient. It helps to establish the ideal disease stage for therapeutic intervention, improves the clinical trial enrolment, and aids in prognostic assessment. In the AD continuum, especially after the FDA approval of anti-amyloid monoclonal antibodies aducanumab and lecanemab there has been an increasing focus on enrolling the key target population of cognitively unimpaired individuals (CU) who are in the preclinical AD stage for clinical trials [7,8]. Visually distinguishing dichotomously negative preclinical AD populations from healthy control for subject selection is challenging. Therefore, detecting and quantifying different protein aggregates from oligomers to fibrils and different aggregate densities from none to frequent can increase the opportunity for potential intervention in early AD cases with disease-modifying therapies.

Particularly for AD, the currently available quantitative metrics for the PET imaging include standardized uptake value ratio (SUVr), z-score, centiloid and the recently reported Aβ load, Aβ index, AMYQ and CenTauR [6,9]. The SUVr, ratio of radiotracer uptake between target and reference region is widely validated against histopathology on a variety of clinical populations. However, the metric is tracer, reference or target region dependent. Additionally, SUVr variability in longitudinal studies has been reported, which limits the accurate detection of time dependent biological differences [10]. The z-score measures the deviation from the cognitively healthy population. The metric is reliant on SUVr values and therefore the weaknesses of SUVr are translated to the z-score. On the other hand, the centiloid (CL) is a universal metric used for inter-tracer standardization of the SUVr through calibration using [^11^C] PiB or a surrogate reference tracer. Recent reports have shown that the magnetic resonance imaging (MRI) is not required for the harmonization of PET image quantification across different tracers using the CL [11,12]. Currently, the CL is mainly validated for global measures and not for the regional uptake. Moreover, the centiloid concordance with the current visual reads is only somewhere in the 25-35 centiloid range [13].

The recently reported Aβ load metric, which is reported in percentage using the [18F] Florbetapir PET tracer, is used for the global estimate of Aβ burden and requires MRI [14]. Initial studies have shown greater sensitivity of Aβ load across all AD classification groups, except in the disease stages prior to the late mild cognitive impairment (lMCI). The Aβ index, another global metric of Aβ pathology, has been reported to show a strong association with PET SUVr. Additionally, it does not require MRI and the definition of regions of interest [15]. The initial reports demand the requirement of universal adaptive template for the direct comparison of values between Aβ tracers without the conversion to centiloid. Similarly, the AMYQ is another metric for global Aβ burden estimate that does not require MRI. It is interchangeable across tracers and does not depend on reference and target regions as indicated by initial cross-sectional study reports [16]. The latest study reports highlighted the requirement of establishing the discriminatory power of these metrics to differentiate patient populations based on disease severity, importantly the distinction between MCI and controls and MCI and AD. On the tau tracer side, CenTauR has been reported for the standardization of quantitative tau PET imaging measures across different tau tracers [9]. However, tau PET tracers are still limited to the clinical experimental studies with only one [18F] Flortaucipir tracer approved by FDA for the imaging of neurofibrillary tangles in cognitively impaired AD patients [17]. Therefore, a detailed validation of other tau tracers is required prior to the evaluation of CenTauR. Additionally, the tracer specific and individual variability in off-target retention needs to be addressed for the harmonization of tau PET images. Overall, the widely validated PET based amyloid quantification methods are global measures and require MRI for spatial registration, priory definition of reference region and the threshold values for positivity are tracer dependent. Additionally, the newly reported metrics, Aβ load, Aβ index and AMYQ are not yet widely validated and unavailable through regulatory approved software.

In the context of fluid-based diagnostic assays for AD, the cerebrospinal fluid (CSF) or plasma biomarker assays rely on cut-off values derived from global estimates of standalone markers or their hybrid ratios. The FDA approved CSF assays for clinical use are hybrid ratios of Aβ 42/40, p-tau 181/Aβ42 and total tau/Aβ42 as shown in Table 1 [18–20]. The positive predictive agreement/ negative predictive agreement, estimated against positive/negative visual reads of PET using FDA approved radiotracers for AD cases, were high. Currently, there are no FDA approved fluid biomarker assays as diagnostic indicators for synucleinopathies. However, recently SYNTap^®^, an α-synuclein seed amplification assay-based CSF test received FDA breakthrough device designation. The assay is yet to be widely validated across different synucleinopathies. Moreover, it is also essential to identify the accurate standard of reference for the estimation of antemortem assay accuracy in addition to clinical diagnosis as comparator. The National Institute on Aging and the Alzheimer’s association (NIA-AA) updated guideline has incorporated the plasma biomarker assays for the diagnosis and staging of Alzheimer’s disease due to its prominent role in recent clinical research [21]. Currently, plasma assays have not received regulatory approval, but a few assays based on standalone amyloid or tau biomarkers have received the FDA breakthrough device designation (Table 1). However, recent studies have shown substantial variation in accuracy of plasma-related A/T/N biomarker assays with respect to PET visual reads or CSF reference standard (60-90%)[22–25].

Recent reports on various therapies targeting earlier stages of protein aggregation including oligomers and protofibrils are exciting. This further underscores the need for reliable biomarker quantification tools capable of accurately detecting and quantifying different forms of protein aggregates even at low levels of burden. Such quantitative biomarker tools are crucial for identifying the eligible patients and monitoring the efficacy of targeted therapies. It is also essential to identify a practical reference standard, beyond the ideal neuropathologic examination, for the validation of tools for the disease diagnosis in the earlier stages. Requirements of radiotracer, post tracer injection time, blood-brain barrier permeability, and off-target tracer binding are the additional challenges associated with the PET imaging. Additionally, the accuracy of the most used PET quantitative metric, SUVr is highly dependent on the reference region and the definition of reference region is challenging especially for longitudinal studies. For example, Moscoso et.al. reported a decreased retention of [18F] Flortaucipir in regions with white matter hyperintensities and up to 57% of the variance in SUVr were linked to the variations in reference region tracer uptake [26]. The recently reported reference region independent PET metrics are for global estimates of biomarkers and the metrics need validation across wider population. The clinically used fluid biomarker assays can also provide the global estimate of biomarker burden without any external contrast agents. However, the global estimates lack the information on spatial distribution of pathology and that makes disease staging challenging. Moreover, regardless of imaging modality or assay, the diagnostic uncertainty exists for values at or near cut-off values for determining normal/abnormal results or on deciding disease staging. In addition to above all, the development of PET tracers for α-synuclein is still an unmet need. Studies have shown that the most accurate and earliest detection of premotor PD should be based on α-synuclein aggregates instead of dopaminergic changes [27]. The lower abundance of α-synuclein, their post-translational modifications, and complex intracerebral environment add challenges to the development of methods for detection and quantification of α-synuclein aggregates. Therefore, it is essential to develop methods for the regional detection and quantification of sparse amounts of biomarkers with different states of aggregation.

Taken together, these considerations motivate the present study, where we propose an X-ray based aggregate index for the regional quantification of protein aggregates in NDDs without a contrast agent and the definition of a reference region. Importantly, the method directly measures the pathophysiologic process by targeting the cross-β structure inherently present in protein aggregates without the requirement of an external contrast agent to indirectly highlight the target. The cross-β structure composed of arrays of β-sheets running parallel to the long axis of the biomarker aggregates and the structure is absent for functional protein monomers [28]. The presence of cross-β rich aggregates were reported previously by X-ray microdiffraction studies for ex-vivo human brain tissues of patients with Lewy bodies of PD [29], amyloid plaques of AD [28], tau deposits of Down syndrome and frontotemporal lobar degeneration (FTD) [30] and glial cytoplasmic inclusions (GCIs) of multiple system atrophy (MSA) [29] (Table S1 in supporting information). The prior studies were limited to the qualitative distinction of healthy and diseased postmortem tissues based on the absence or presence of X-ray cross-β signals of matured biomarker deposits. The current study addresses the potential of X-ray scattering technique to detect and quantify cross-β signals of oligomeric and fibrillar deposits in tissues from different brain regions without the requirement of a reference region. Additionally, we demonstrate a material decomposition method using sheep brain model and phantoms for the quantification of regional protein aggregates based on cross-β signals deconvolved from complex heterogeneous diseased tissues. The proposed method shows the applicability for both synchrotron as well as laboratory-based X-ray sources to further expand the scope of this study.

## 2. MATERIALS AND METHODS

The seven months old sheep head was purchased from a local butcher shop. Skull and dura mater were removed prior to the brain sectioning. The brain was sliced into two at the mid-sagittal plane. Each slice was further sectioned into ten pieces and the details of the tissue regions are discussed in Section 3.5 of this article. Brain samples were stored in the freezer (temperature ∼ -15 °C). Prior to the experiments, tissues were thawed in the refrigerator (temperature ∼ 4 °C). Tissues were sealed in the zip-lock cover and those were at room temperature for about ten minutes during the experiments.

The bovine serum albumin (material number: A7030, BSA), β-lactoglobulin from bovine milk (lyophilized powder, material number: L3908, genetic A and B mixture, LGAB) and hydrochloric acid (1.0 N solution, material number: H9892, HCl) were purchased from Sigma–Aldrich (St. Louis, MO). The fat (Sweet cream salted butter with pasteurized cream and salt) was purchased from a local grocery shop (Giant Food of Maryland, LLC). All chemicals were used as received without further purification.

### 2.1. Preparation of synthetic oligomer and fibril

As received bovine serum albumin (BSA) was used as the model system for the pathogenic oligomer β-sheets. Fibrils of β-lactoglobulin were used as the model system for pathogenic fibrillar β-sheets. Fibrillization protocol and detailed characterization of these oligomer and fibrillar model systems were reported previously [31,32]. Briefly, LGAB fibrils were prepared by incubating the 3 wt.%, pH 2 LGAB solutions in a 90 ^0^C water bath for 12 h followed by rapid quenching of the solution in an ice bath for 1 h. The pH 2 solution of LGAB was prepared by adjusting the pH of aqueous LGAB solution to 2 by adding 1.0 N HCl and by accurately monitoring using a pH meter (Mettler Toledo, FiveEasy). The protein secondary confirmation and nanoscopic structure of oligomer and fibrils were characterized in detail using Attenuated total reflection Fourier transform infrared (ATR-FTIR) spectroscopy, solution small angle X-ray scattering, monochromatic and polychromatic X-ray scattering at wide angle and the details are reported elsewhere [32].

### 2.2. Low energy (8 keV), monochromatic wide-angle X-ray scattering (WAXS)

Low energy (8 keV), monochromatic wide angle X-ray scattering (wave vector, 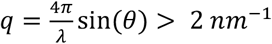, where 2*θ* is the scattering angle, and wavelength, *λ* = 0.154 *nm*) experiments were performed using a SAXSpace system (Anton Paar, Graz, Austria). The instrument was equipped with a conventional X-ray generator (Seifert Analytical X-ray, 40 kV, 50 mA), copper-K_α_ (Cu-K_α_) anode, micro-strip photon counting X-ray detector (Mythen, Dectris Ltd., Baden, Switzerland) and a water chiller (Haskris, IL, USA). The sample to detector distance (SDD) was 121 mm and the scattering vector was calibrated using silver behenate. The irradiation beam was line collimated, and a semitransparent beam stop attenuated the highly intense primary X-ray beam. The primary beam is used to determine the zero-angle or zero-wavevector (*q*_0_,) position. The relative intensity of scattering data was further corrected using the transmittance of the direct X-ray beam through sample followed by background subtraction. WAXS data were linearly binned to 100 bins and de-smeared using Lake algorithm [33]. The above explained standard corrections were performed using Anton Paar SAXSanalysis software (Austria, version-4.20.048). Finally, the 1D data was baseline corrected using a fourth order polynomial fitting between *q* = 3 *nm*^−1^ *to* 18.8 *nm*^−1^.

For WAXS studies, samples were packed in adhesive tape and sealed within the rectangular slit shaped sample holder (slit width 2.8 mm and length 20 mm) to ensure the whole sample is within the incident X-ray path and exposed to X-rays (Figure S1 in supporting information). The butter-oligomer phantoms were prepared by stacking as two layers where butter at the bottom and oligomer model were stacked by compressing on top of that as shown in Figure S1. Background correction was performed by subtracting the scattering from adhesive tape and background scattering signal was collected at the same experimental conditions as the sample. Total X-ray exposure time for each sample was 1 h (combined 4 frames where each frame was taken for 900s X-ray exposure). Minimum three independent measurements were performed using freshly prepared phantoms for each formulation and the standard errors were estimated as the deviation of values from their means.

### 2.3. High energy (30-80 keV), polychromatic spectral X-ray scattering (sXS)

The instrument technical details can be found in the previous publications [34,35]. Briefly, sXS is comprised of high energy, polychromatic X-ray source (tungsten anode, MXR-160/22, COMET), two pinhole collimations, lead beamstop with a 300 µm hole and spectroscopic photon counting detector (HEXITEC, Quantum Detectors, Oxfordshire, UK). Polychromatic X-rays were generated using 80 kVp tube voltage and 1 mA current and X-ray exposure time of 600 s unless otherwise stated. Two pin hole lead collimators were used for the X-ray beam collimation. The first pin hole (P1) diameter was about 5 mm and the 2 mm diameter second pin hole (P2) was placed 215 mm away from P1. The sample to detector distance (SDD) was 460 mm and the X-ray path between source and the detector was non-vacuumed. Cadmium telluride (CdTe) energy resolving photon counting pixelated detector was 1 mm thick and it had 80 × 80 pixels with 250 µm pixel pitch. The scattered photons were collected between 30-80 keV energy range using 27 keV energy threshold and 1 keV energy binning. For sXS, the photon energy (*E*) and scattered angle (*θ*) dependent wavevector (*q*(*E, θ*)) was calculated based on the equation:

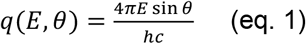

where, *hc*=1.24 eV-µm. The charge sharing correction, energy dependent transmission correction followed by summing of corrected *q* data for all energy bins were performed as reported before. Finally, the background scattering was subtracted from the sample scattering profile. In this study the background scattering was from adhesive tape alone or from both adhesive tape and Ziplock® cover. The background measurements were performed under the same experimental conditions as sample. The instrument was calibrated using caffeine powder.

The model oligomer and fibril samples were wrapped in adhesive tapes as pouches where the samples were localized in a 2 mm^2^ area. Protein sample pouches were directly adhered on the second collimator (P2) with 2 mm hole diameter and prior to every measurement, we ensured that the pouches are within the collimator hole. For sheep brain experiments, the brain tissues sliced from different brain regions were packed inside the thin Ziplock® covers and those were directly adhered either to the P2 or on top of the protein sample pouches depending on the experimental conditions. For each brain samples, the region of interest (ROI) was marked, and the X-ray was shot within the ROI for each measurement. The reported tissue thicknesses were the average of thicknesses measured within the ROI.

### 2.4. Analysis pipeline for material decomposition and component quantification from X-ray spectrum

The processing of X-ray spectrum involves signal decomposition and deconvolution. Spectrum decomposition involves decomposing signal into sum of individual components that characterize the properties of different constituents in a material. This approach improves the separability of overlapped peaks including the extraction of low amplitude X-ray signals (weak echoes). In our study, we used Gaussian decomposition with an assumption that the shape of the scattered signal is Gaussian-like and the received signal is regarded as the convolution of the scattered signals of the differential scatters. The Gaussian decomposition involves the optimization of parameters that define the component peaks. The analysis of X-ray scattering spectrum invariably involves some or all the steps shown in the process pipeline (Figure 1). Peak selection and annotation involve the identification of number of merged peaks and their peak locations followed by annotation of each peak to the nanoscopic structural characteristics of X-ray scatters present in the material. The structural information of each material constituents was reported using the following peak parameters and the listed metrics were used for the evaluation and comparison of material compositions:

**Figure 1:**
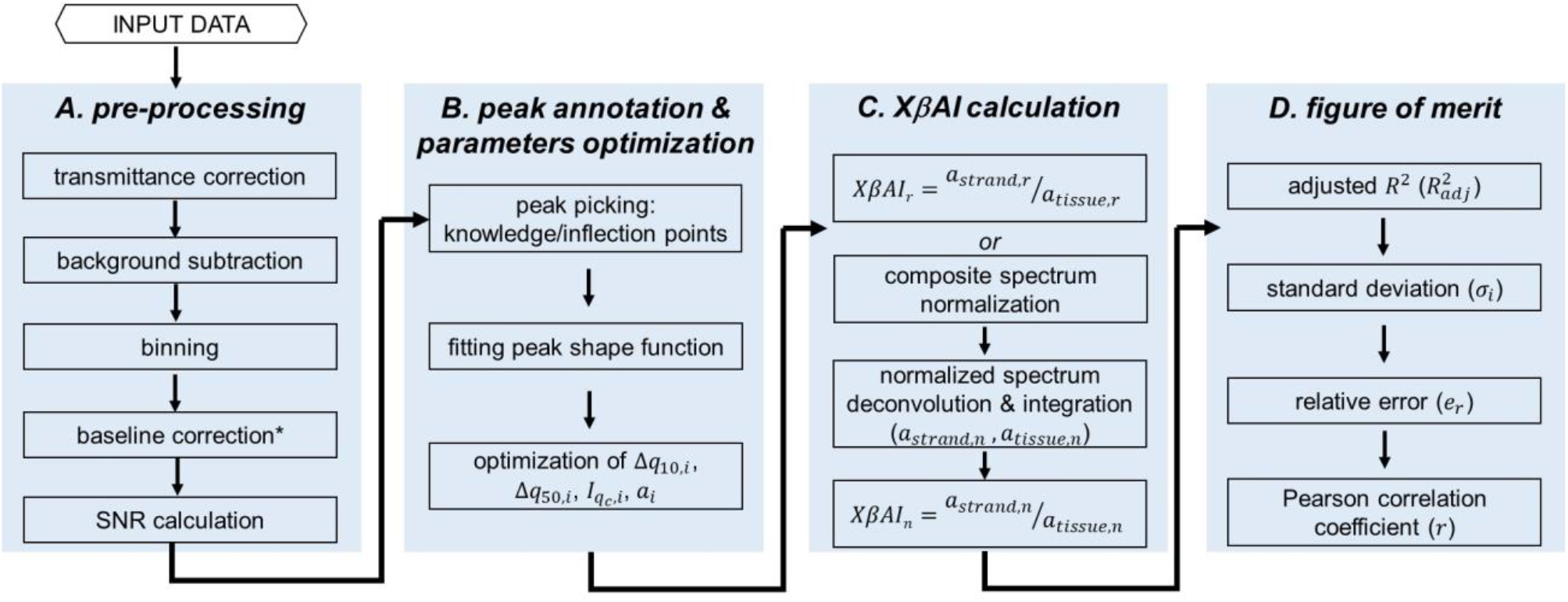
Flow chart of X-ray spectrum analysis pipeline for the material decomposition, quantification of material compositions, estimation of XβAI and the figure of merits used for evaluating the performance of spectrum analysis protocol. ^*^The baseline correction was performed only for the WAXS signals collected using monochromatic, low energy X-ray source.

### i. Fixed parameters

a. Total peak numbers and peak locations (*q*_*c,i*_ for component *i*): The knowledge of expected total number of scatter components and their peak positions were exploited in our peak-searching program. The initial parameters of the Gaussian model are estimated through inflection points from the deconvolved waveform. Annotation of peak locations are either knowledge based or a combination of prior knowledge and local maxima of shoulder peaks or inflection points in the second derivatives of the composite signal. There are also cases where the inflection points are not evident for weak echoes overlapped in a composite signal either due to low concentration or weak scattering of constituents. In such cases, the peak locations are assigned based on the prior knowledge of the material composition and their corresponding X-ray scattering peak positions.
b. Peak function: Gaussian peak shape models are used to fit the overlapped peaks of individual components in a composite spectrum. The validity of Gaussian function was validated either based on asymmetry factor 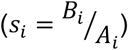 or peak width ratios.

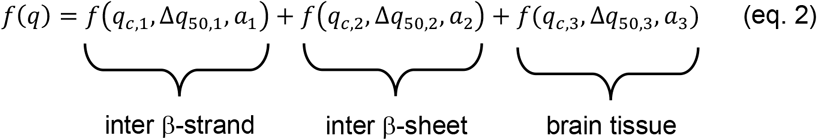

where, 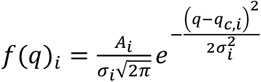 for a Gaussian function and 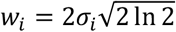

### ii. Floating parameters

a. Left (*A*_*i*_) and right (*B*_*i*_) widths at 1/10 the maximum ordinate of the peak
b. Full width tenth maximum (FWTM, Δ _*q* 10,*i*_): distribution width at a level that is 1/10 the maximum ordinate of the peak
c. Full width half maximum (FWHM, Δ _*q* 50,*i*_): distribution width at a level that is 1/2 the maximum ordinate of the peak
d. Peak amplitude 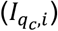 : peak height at peak location *q*_*c,i*_
e. Peak area, 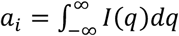 The area under the peak for each peak is used to estimate relative quantities of each component in a material.

The least square method and optimization algorithm were used to determine the floating parameters that best fit the decomposition profile. It was ensured that the signal to noise ratio (SNR) of composite signals are high and the minimum peak values were greater than the noise threshold. The noise of X-ray signal was extracted by performing the Daubechies wavelet denoising as described reported before [36]. The trend and detrend signals were extracted by fitting the experimental X-ray scattering signal, *f*(*x*) to the Daub4 basis set.

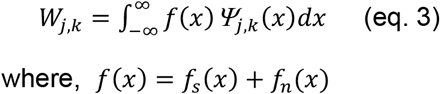

Here, *Ψ* (*x*) is the wavelet function, *f*_*s*_(*x*) and *f*_*n*_(*x*) are the trend and detrend signals of the input X-ray signal *f*(*x*), *j* and *k* are the scale and displacements.

The SNR was calculated as the

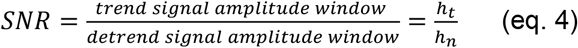

The peaks that do not meet the conditions were deleted and re-performed the optimization until the final solution is obtained. Accuracy of estimated values by Gaussian material decomposition were assessed by comparing with the ground truth.

### 2.5. Figures of merit

The reliability and repeatability of each measure were ensured by calculating the following metrics under various realizations.

#### 1. Adjusted coefficient of determination for goodness of fit

The coefficient of determination (*R*^2^) measures the overall quality of the regression as it is the percentage of total variation exhibited in the data that is accounted for or predicted by the regression line.

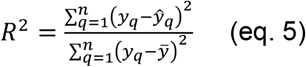

where, *ŷ*_*q*_ is the regression estimate and 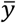 is the *y* mean.

Adjusted 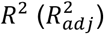 accounts for the number of variables (*k*) and the sample size (*n*) and modifies *R*^2^ as:

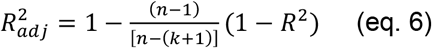

The rule of thumb followed for the peak shape function fitting was *n* ≥ 5(*k* + 2).

### 2. Standard deviation (*σ*_*i*_)

The standard deviation indicates the spread of quantities of a component *i*, derived from multiple measurements (*N*) under similar acquisition conditions.

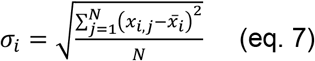

where, 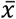 is the mean value.

### 3. Relative error (*e*_*r*_)

The relative error is used to represent the deviation of a quantity (*x*_*i*_) derived from a composite spectrum from its standard of reference (*T*) obtained under similar acquisition conditions.

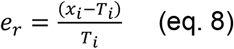

### 4. Pearson correlation coefficient (*r*)

This represents the degree of association between decomposed quantity from a composite spectrum with its corresponding standard of reference or ground truth.

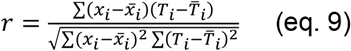

where, 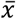 and 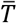 are the mean values.

## 3. RESULTS

The decomposition and quantification of healthy or diseased brain tissue material to extract relative contribution of each component was performed by knowledge based peak decomposition method. This method is evaluated for different categories of experimental data:

1. Broad high *q* range (*q* = 4.6 *nm*^−1^ *to* 45 *nm*^−1^) WAXS spectra of healthy human brain white matter and grey matter
2. Brain tissue phantoms with known proportions of fat and synthetic protein oligomer model aggregates
3. Varying clinically relevant quantities of synthetic model aggregates of oligomers and fibrils
4. Sheep brain tissue with known quantities of synthetic aggregate models of oligomers and fibrils under various realizations. The realizations are:
  a. Fixed oligomer loading at different brain tissue regions.
  b. Varying oligomer loading at middle temporal brain tissue region.
  c. Varying oligomer loading at olfactory bulb brain tissue region.
  d. Varying fibrillar loading at frontal cortex tissue region.

Finally, we establish a quantitative metric, the X-ray cross-β aggregate index (XβAI), for the representation of regional protein aggregate density in different tissue regions.

### 3.1. Material decomposition of human brain white matter and grey matter tissues

The initial demonstration of the proposed material decomposition approach was performed on the WAXS profiles of human brain white and grey matter (Figure 2). The WAXS (synchrotron X-ray source, ID02 beamline/ESRF, 12.4 keV, SDD-1.2 m) profiles of grey and white matters were adopted from reference [37]. The grey and white cerebral matter samples were from 90- and 60-year-old female and male respectively. The tissues were not known for any neuropathological disorders. Both, grey matter and white matter exhibited three peaks with peak locations (*q*_*c*_) around 14.2 nm^-1^, 19 nm^-1^ and 27 nm^-1^ (Figure 2-b,c, Figure S2 in supporting information). The peak at the 14.2 nm^-1^ is due to the lipid component of the tissue and peaks at 19 nm^-1^ and 27 nm^-1^ are arising from aqueous component. The cumulative fits with 3 peaks decomposition using Gaussian function yielded 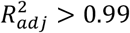. The quantitative assessment by each peak integration yielded, ∼72% water and ∼28% total lipid for white matter (Figure 2-b). Whereas the lipid fraction is lower for grey matter (∼11%) and about 89% water for grey matter (Figure 2-c). The fat and water proportions quantified from X-ray scattering spectrum were comparable with the literature [38]. The reported aqueous component is between 80-85% for grey matter and 75-80% for white matter. Therefore, the material decomposition and their quantification using peak shape analysis approach was successful for postmortem healthy human brain tissues.

**Figure 2:**
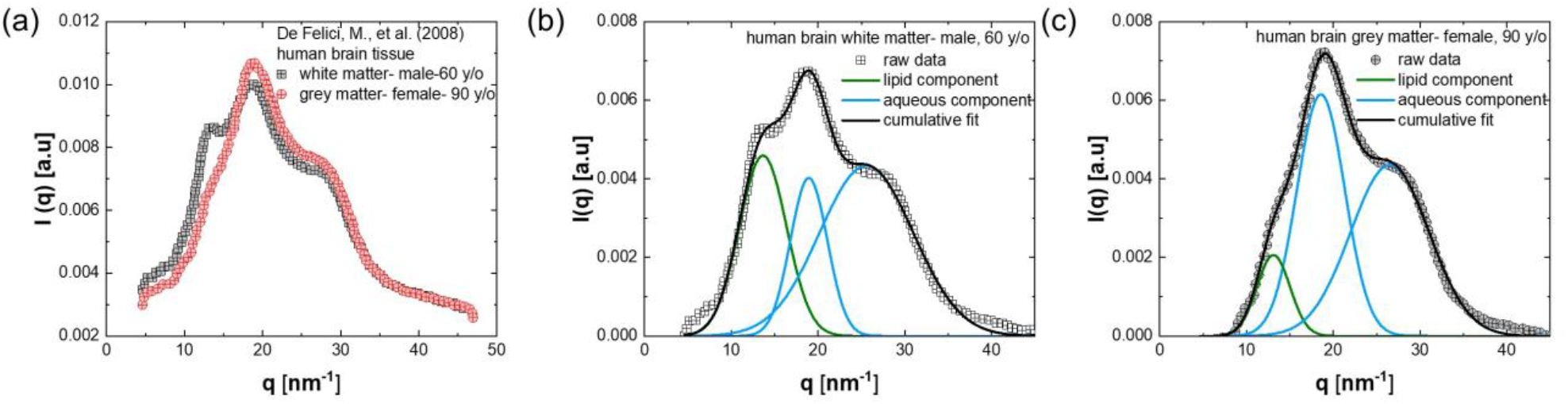
(a) Synchrotron X-ray source based WAXS profiles of human brain white and grey matter tissues. The data was adopted with the permission from reference [37]. The spectra of (b) white matter and (c) grey matter were decomposed using Gaussian function to three broad peaks centered on q_c_ =14.2 nm^-1^ (green line), 19 nm^-1^ and 27 nm^-1^ (both blue lines) are peculiar of tissue composition, where the first peak is characteristic of the lipid content and the other two are due to the aqueous component.

### 3.2. Brain tissue phantoms with known proportions of fat and synthetic oligomer model aggregates

Next, we prepared brain tissue phantoms with known amounts of oligomer model aggregates. The bovine serum albumin with about 18% β-sheet content and antiparallel cross-β motifs was used as the oligomer model aggregate. We have previously reported in detail the preparation, characteristics of model protein aggregates and their property comparisons with aggregates of amyloid β-42 and α-synuclein [32]. The oligomeric and fibrillar models for amyloid β-42 and α-synuclein were established by considering the position of the β-sheet peak in the amide I infrared absorption band, the percentage of β-sheet content, the oligomer index, the size and shape of aggregates at the nanometer length scale, and the peak positions in the wide-angle X-ray scattering spectrum (for detailed comparisons between the model aggregates and aggregates of Aβ-42 and α-synuclein, refer to [32]). The model aggregates were stacked by compressing on top of tissue phantom material (see section 2 for further details). The weight proportions between tissue phantom material and oligomer model were known for each formulation (Table S2). During the WAXS experiments, it was carefully ensured that the whole material is exposed to incoming X-ray beam (Figure S1). The WAXS spectrum of oligomer model (*weight, w* = 7 *mg*) exhibited two broader peaks with peak locations around 6.4±0.2 nm^-1^ (*q*_*c,sheet*_) and 13.5±0.3 nm^-1^ (*q*_*c,strand*_) (Figures 3a, S3a in supporting information). The X-ray scattering spectrum was similar to the previously reported cross-β antiparallel profile of anisotropic oligomer aggregates of amyloid-β in AD patients [39]. The peaks correspond to the inter-β-sheet (*q*_*c,sheet*_) and inter-β-strand (*q*_*c,strand*_) spacings, as well as to arrays of β-sheets running parallel to the long axis of the aggregate (Figure 3b, inset). The peak decomposition of tissue phantom X-ray spectrum showed three peaks with peaks centered around 14.3±0.3 nm^-1^, 15.9±0.2 nm^-1^ and 17.5±0.4 nm^-1^ (Figure 3a and S3b in supporting information). The *q* range for these measurements were limited between 3-to 18.7-nm^-1^. Therefore, the aqueous component peak was either at the tail of the baseline corrected spectrum (*q*_*c*_ = ∼ 18 *nm*^−1^) or at the second peak (*q*_*c*_∼ 27 *nm*^−1^) observed for healthy human brain white and grey matter tissues in Figure 2. However, this second peak is not accessible in these measurements. The deconvolution of composite spectrum of tissue phantom embedded with oligomer exhibited five peaks. In a blend with 7 mg oligomer and 28 mg tissue phantom 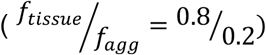, the peak locations were at *q*_*c*_ = 6.7 *nm*^−1^, 13.3 *nm*^−1^, 14.4 *nm*^−1^ 15.9 *nm*^−1^, *and* 17.4 *nm*^−1^. These peaks originated from both tissue phantom and oligomer model (Figure 3b). The oligomer peaks became more compact in blends compared to standalone formulation, which could be due to packing of these aggregates within the phantom tissue matrix. The ratios of area under the peak (AUP) of deconvoluted peaks of inter-β-strand and inter-β-sheet spacings to the total AUP of composite spectrum were 0.198±0.1 and 0.11±0.1 against ground truth value of 0.2 for 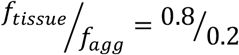 formulation. The data suggests that the AUP of inter-β-strand spacing quantitatively represent the total aggregate loading in the blend. We further validated the finding by analyzing other formulations reported in Table S2. Here we varied the oligomer loading from 0 to 7 mg which corresponds to *f*_*agg*_ from 0.003 to 0.2. In this case, increase in oligomer loading increased the composite spectrum width compared to its amplitude (Figure 3c). The measured AUP ratios of inter-β-strand (*f*_*agg,meas*_) were comparable with the known fractions of aggregates (*f*_*agg,true*_) in each formulation (Figure 3d, Table S2). The plot between the measured aggregate fractions (*f*_*agg,meas*_) and the corresponding ground truths (*f*_*agg,true*_) were strongly correlated with a Pearson correlation coefficient greater than 0.98. The relative error for each fraction was less than 0.02. This study confirms that the material decomposition based on peak shape analysis on composite X-ray spectrum can accurately quantify even very small oligomer aggregate fractions (*f*_*agg*_ = 0.003) from the tissue mimicking phantom environments. For further studies, the aggregate burden will be quantified as the area under the peak of inter-β-strand spacing with *q*_*c,strand*_ ∼13.3 *nm*^−1^.

**Figure 3:**
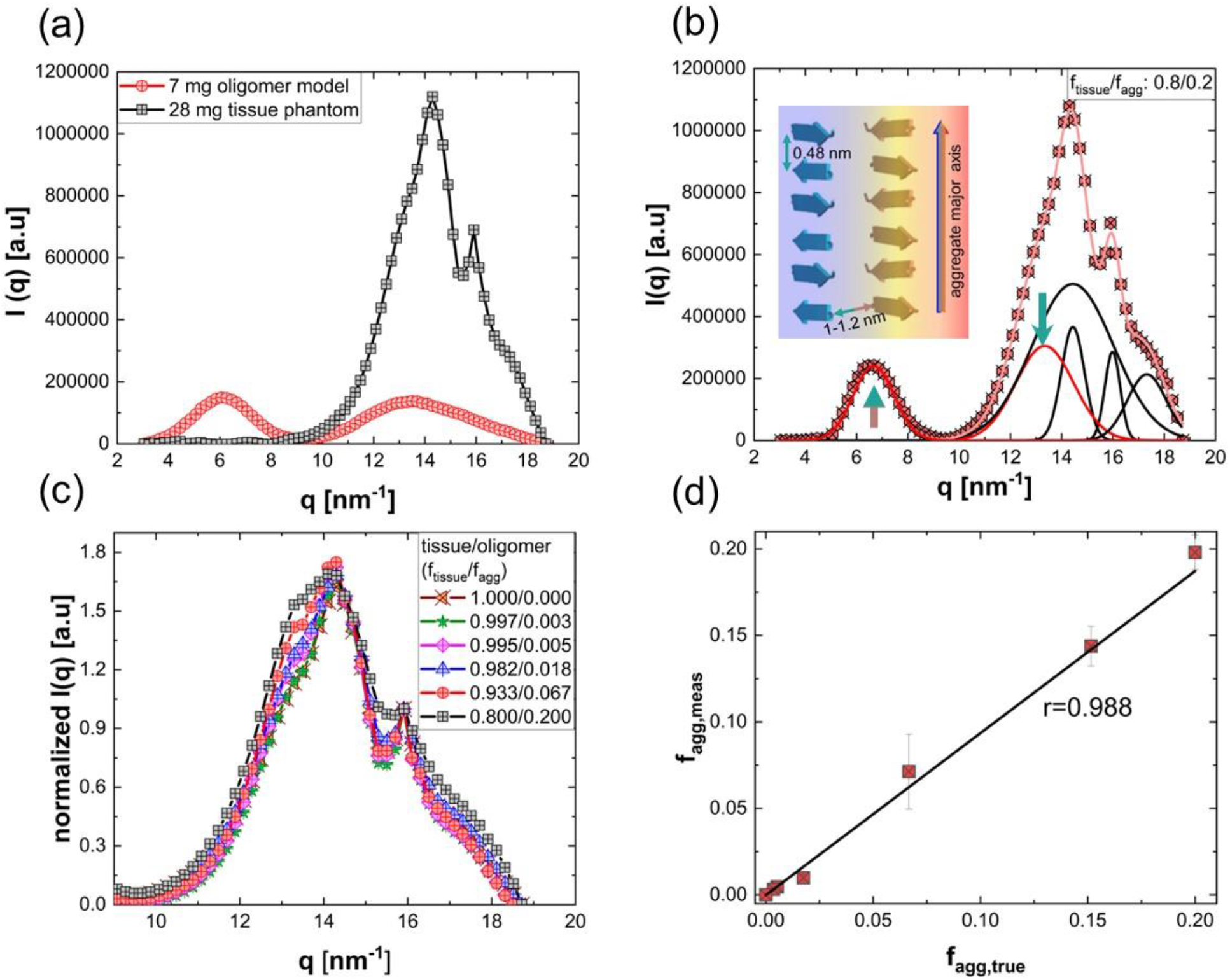
The monochromatic X-ray source based WAXS profiles of brain tissue phantom, model oligomer aggregates and their blends with different proportions of fat and oligomer. (a) Individual WAXS profiles of 7 mg oligomer and 28 mg tissue phantom. (b) WAXS profile of a blend made by mixing 7 mg oligomer to 28 mg tissue phantom (weight proportions between fat and oligomer is,f_tissue,true_: f_agg,true_ = 0.8: 0.2). The pre-processed experimental data is shown using symbols and the decomposed oligomer and tissue profiles are shown using red and black line curves respectively. The inset schematic shows the arrangements of β-strands and β-sheets in the cross-β substructure. The inter-β-sheet spacings and β-strand spacings are illustrated with colored arrows, and the corresponding WAXS peaks associated with these spacings are indicated using arrows of the same color for straightforward interpretations. The simulated cumulative fit is shown with light red line curve and the goodness of fit, 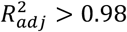. (c) WAXS profiles of blends prepared with different proportions of tissue phantom and oligomer. The profiles were normalized with the X-ray intensity at q_c_ = 15.9 nm^−1^ to homogenize the tissue profiles and to highlight the effect of oligomer quantity on composite profile. (d) The plot shows the concordance between true aggregate fraction (f_agg,true_) known during sample preparation and the aggregate fraction (f_agg,meas_) estimated by integrating the decomposed peak of inter-β strand spacing. The error bar is the estimated relative error.

### 3.3. X-ray scattering signals of clinically relevant quantities of synthetic model aggregates of oligomers and fibrils

We further studied the X-ray scattering signals of tissues from different brain regions with and without protein aggregates. In this study, we opted for the sheep brain as our model due to its comparable neuroanatomical and neurofunctional features with the human brain, emphasizing its suitability for translational research. Clinically relevant quantities of model oligomers and fibrils were embedded in the different anatomical regions of the brain. We then evaluated the accuracy of quantitatively decomposing X-ray signals from both normal and artificially diseased tissues. In this context, to establish the ground truths, we first individually studied the X-ray scattering properties of tissues from different brain regions and clinically meaningful quantities (50 µg to 6 mg) of model oligomers and fibrils. Given that brain tissues exceed a thickness of 1 mm, we conducted WAXS studies utilizing spectral X-ray scattering techniques. This involved employing a polychromatic X-ray source in conjunction with a photon-counting detector.

The oligomer model under investigation comprised BSA aggregates with a β-sheet content of 18%, characterized by antiparallel β sheets. The quantities of this oligomer were systematically varied, ranging from 50 µg to 6 mg (Figure 4a). The aggregates were wrapped into 2 mm size spheres in adhesive tapes and adhered onto a 2 mm diameter pin hole to ensure the major portion of sample is interacting with the incoming X-ray beam. Nevertheless, ensuring the complete exposure of the entire sample to X-rays posed a challenge. Intriguingly, X-ray signals were detected even for 50 µg oligomers utilizing a laboratory X-ray source, operating under experimental conditions of an 80 kVp tube voltage, 1 mA current, and a 600s X-ray exposure time. As the aggregate amount decreased, there was a corresponding decrease in the signal-to-noise ratio (SNR). The SNR increased with increase in X-ray exposure time (Figure S4a). The increase in current from 1 mA to 5 mA for a fixed voltage and X-ray exposure time did not show a noticeable effect on the scattering profile (Figure S4b). The X-ray signals were broad for all oligomer quantities, and it was challenging to assign peak locations for inter β-strand (*q*_*c,strand*_) and inter β-sheet (*q*_*c,sheet*_) spacings based on local peak maxima or second derivatives due to the presence of noise (Figure S5). Hence, *q*_*c,strand*_ = 13.3 *nm*^−1^ and *q*_*c,sheet*_ = 6.5 *nm*^−1^ were established based on pre-existing knowledge and the β-strand and β-sheet peaks were decomposed from the composite profiles of each oligomer quantity. The area under the peak of the decomposed inter-strand spacing signals 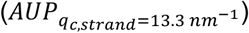 exhibited an increase with the increase in oligomer quantity, showing a scaling exponent of *n*∼0.24 (Figure 4b).

**Figure 4:**
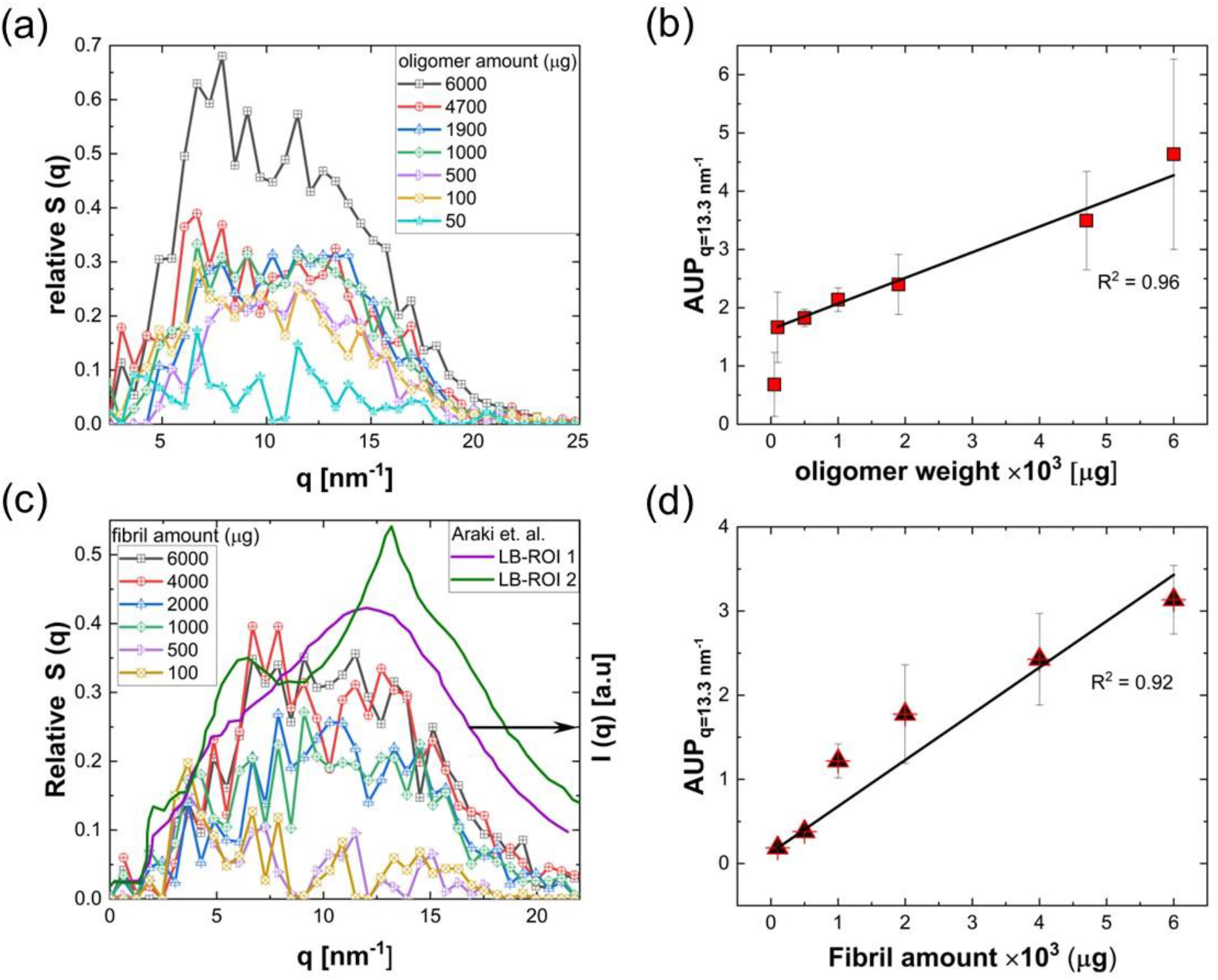
The WAXS profiles of clinically relevant quantities of model oligomers and fibrils collected using spectral X-ray scattering (sXS). Fibrillar spectra comparison with Parkinson’s disease Lewy bodies X-ray diffraction spectra. (a) Scattering profiles of BSA oligomers (%β sheet: 18, antiparallel β sheets) with quantities ranging from 50 µg to 6 mg. The relative s(q) is the scattering cross-section normalized to the primary X-ray beam and background subtracted. The signal preprocessing details are described in section 2.3. The scattering profiles were decomposed to two peaks with peak locations, q_c,strand_ = 13.3 nm^−1^ and q_c,sheet_ = 6.5 nm^−1^ and calculated the area under the peak (AUP) of decomposed profiles by integration. (b) AUP of decomposed peak with q_c,strand_ = 13.3 nm^−1^ as a function of oligomer amount shows an increase with scaling exponent of n∼0.24. (c) WAXS profiles of fibrillar model, LGAB fibrils (%β sheet: 55, parallel β sheets), with varying quantities from 100 µg to 6 mg. The synchrotron X-ray source based WAXS spectra of Lewy bodies from age matched, two different PD patients are shown for the comparison. The synchrotron data is adopted with the permission from [29] (d) AUP of decomposed inter-β strand peaks of fibrils with q_c,strand_ = 13.3 nm^−1^ as a function of fibril amount shows an increase with the scaling exponent of n∼0.72. The coefficient of determination (R^2^) is shown for each fit.

The X-ray scattering properties of the fibrillar model, LGAB fibrils (%β sheet: 55, parallel β sheets), were also studied by varying the quantity from 100 µg to 6 mg (Figure 4c). The photon count increased linearly with the fibril amount (Figure 4d). In this case as well, the inter-β strand and inter-β sheet peaks were not distinctly separated (Figures 4c and S6). The X-ray profiles of the model fibrils were compared with the previously reported X-ray signals of Lewy bodies (LBs) formed from α-synuclein aggregates in Parkinson’s disease patients (Figure 4c, line profiles). Previous findings have demonstrated diversity in the X-ray signals of α-synuclein aggregates both within a patient and among age and sex-matched neuropathologically confirmed PD patients (Figure S6 and Table S1 in supporting information) [29]. The isolated peaks typical of amyloid fibrils with locations at *q*_*c*_∼6.1 and 13.5 *nm*^−1^ were observed in some LBs, for instance, in LB-ROI 2 as shown in Figure 4c. The X-ray microdiffraction profiles of LB-ROI 1 and LB-ROI 2 were reported by Araki et. al. and the data were collected using synchrotron micro-X-ray beam at the BL40XU beamline [29]. However, similar to the X-ray spectrum of model fibrils, broader X-ray spectra without distinct peaks were also observed in Lewy bodies, as illustrated by LB-ROI 1 in Figure 4c. The diversity in the X-ray spectra of specific protein aggregates from age and sex-matched patients, or within the same patients, could arise from differences in the maturity and stacking regularity of β-sheets. Importantly, our comprehensive qualitative and material decomposition-based quantitative analyses of previously reported X-ray spectra from aggregates in ex-vivo brain tissues of patients with various neurodegenerative disease, animal models, and recombinant protein aggregates confirm the diversity in the X-ray signals of cross-β motifs (Table S1 and Figure S6 in supporting information). To the best of our understanding, this is the first detailed analysis of cross-β motifs in varying biomarkers associated with different NDDs. Our analyses also suggest that pre-defining peak shape characteristics, such as width, height, and sharpness, is impossible even for the same type of protein aggregates. The increase in fibril inter β-strand 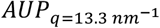 with fibril amount was steeper than oligomers with a scaling exponent, *n*∼0.72. Interestingly, the ratios of scaling exponents for oligomers and fibrils matched the ratios of their % β-sheet content in those aggregates 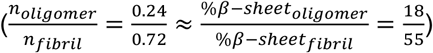. This further confirms that the X-ray signals solely originate from the β-sheets in the protein aggregates.

### 3.4. X-ray signals of sheep brain tissues with varying fibrillar and oligomer loadings

### 3.4.1. Varying fibrillar amount in frontal cortex

The amount of fibrillar model systematically varied from 100 µg to 6 mg in a region of interest chosen within the frontal cortex tissue (Figures 5a and S7). The X-ray spectrum of tissue was centered around *q*_*c,tissue*_ ∼12.6 *nm*^−1^. The addition of 100 µg fibril to the tissue led to about 8% increase in X-ray spectrum amplitude and full width half maxima (Δ*q*_50_). The change in spectrum width with the addition of fibril was not evident for full width tenth maximum (FWTM, Δ*q*_10_). Further increase in fibril amount systematically increased the spectrum amplitude (Figure S8a), whereas the Δ*q*_50_ did not show significant change with further increase in fibril amount from 100 µg (Figure S8b). The Δ*q*_10_ did not show any specific trend with an increase in fibril loading (Figure S8c). The X-ray signals of tissue, fibril inter β-sheets and fibril inter β-strands were decomposed from the composite X-ray spectrum using the material decomposition pipeline discussed in section 2.4. Here, the fibril signals have low echo compared to the tissue signals as the fibril quantities are lower compared to the tissue amount. Therefore, the inter β-sheet and inter β-strand signals are hidden within the composite spectrum and their peak locations cannot be obtained from spectrum second derivatives. For the material decomposition, we have used the prior knowledge of peak locations. To confirm the accuracy of decomposition approach, the deconvoluted signals of each component were compared with their ground truths. Here, the ground truths were the X-ray signals collected for individual components with matched -quantities and -X-ray exposure locations. For example, the signal decomposition of composite spectrum collected from frontal cortex tissue with 4 mg fibrils is demonstrated in Figure 5b. The peak locations of decomposed signals were around *q*_*c,tissue*_ ∼12.7 *nm*^−1^, *q*_*c,strand*_∼13.35 *nm*^−1^ and *q*_*c,sheet*_∼6.74 *nm*^−1^. The decomposed tissue signal (black line curve in Figure 5b) matched very well with the tissue alone signal acquired separately. The cumulative fit of decomposed inter-β-sheet and inter β-strand signals matched well with the 4 mg fibril signal alone and with signal obtained by subtracting the signals from tissue alone and tissue with 4 mg fibrils (Figure 5b). The area under the peaks (AUPs) of decomposed inter β-strand signals were compared with their reference standard (Figure 5e). The AUPs were very well correlated and the Pearson’s correlation coefficient (*r*) was around 0.98.

**Figure 5:**
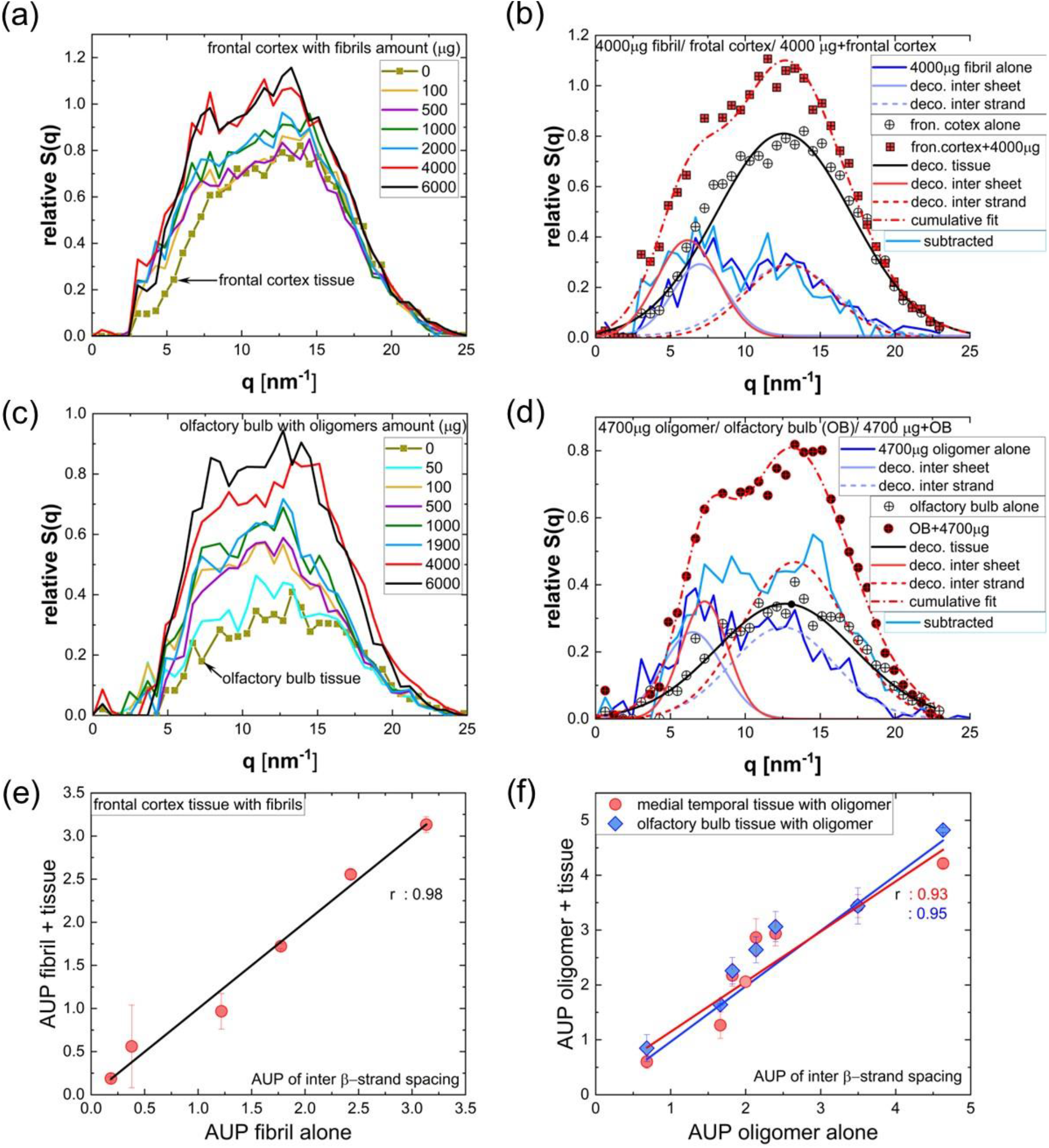
The WAXS profiles and their characteristics of sheep brain frontal tissue with varying fibrillar loadings and olfactory bulb tissue with varying oligomer are illustrated in Figure (a). The WAXS profiles depict the frontal cortex tissue both before and after the addition of different quantities of LGAB fibrils. (b) The figure demonstrates the material decomposition process based on the knowledge of peak locations for frontal cortex tissue with a fibril loading of 4000 µg (4 mg). The decomposed inter-β-strand signal is represented by the red short dash line, while the inter-β-sheet signal is shown by the red solid line, both derived from the composite spectrum (rectangular symbol with a plus interior). These decomposed signals are then compared with the signals obtained from the 4 mg fibril alone (light blue solid and short dash lines). Additionally, the tissue signal decomposed from the composite spectrum (black solid line) is compared with the ROI-matched frontal cortex tissue alone signal (black circle and plus interior). The composite spectrum subtracted from the ROI-matched tissue alone signal is indicated as the ‘subtracted’ signal (shown with a shade of cyan blue color line) and is compared with the 4 mg fibril alone signal (blue line). (c) WAXS profiles of olfactory bulb with varying oligomer quantities. (d) The illustration showcases the material decomposition process from the composite spectrum for the olfactory bulb with a 4700 µg (4.7 mg) oligomer loading. The decomposed signals are then compared with the ROI- and quantity-matched tissue and fibril alone signals, similar to the demonstration in Figure 5b. The composite signal subtracted from the ROI-matched tissue alone signal is compared with the quantity-matched oligomer alone signal. The area under the peak of inter β-strand signal decomposed from composite spectrum of (e) frontal cortex tissue with fibrils and (f) olfactory bulb and tissue from medial temporal lobe with oligomers as a function of fibrillar and oligomer burden respectively.

### 3.4.2. Varying oligomer amount in olfactory bulb and medial temporal lobe tissues

The amount of oligomer model within the tissue was systematically varied from 50 µg to 6 mg (Figures 5c and S9). The tissue regions of interest were olfactory bulb and medial temporal lobe. The X-ray spectra of both tissues were centered around *q*_*c,tissue*_ ∼12.7 ± 1 *nm*^−1^. Similar to the case of fibrils in the frontal cortex, the spectrum amplitude increased with the increase in oligomer loading to the olfactory bulb and medial temporal lobe (Figures S10 and S11 in supporting information). The addition of 50 µg oligomer to the olfactory bulb led to about 12% increase in X-ray spectrum maximum amplitude and about 36% increase with the addition of 100 µg oligomer (Figure S10a). The increase in maximum spectrum amplitude was about 5- and 11-% for medial temporal lobe tissue with 50- and 100-µg oligomer additions respectively (Figure S11a). For a fixed oligomer volume, the X-ray interacting tissue region volumes (*V*_*tissue*_) were 3.5 mm^3^ and 9.4 mm^3^ for olfactory bulb (1 mm X-ray beam radius, 1.1 mm tissue thickness) and medial temporal (3mm thickness) tissues respectively. The variation in the percentage increment in composite spectrum amplitude between the tissues of the olfactory bulb and the medial temporal lobe after the addition of oligomers may be attributed to differences in the proportions of volumes between oligomer and the tissue region 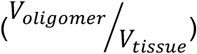. The spectrum width did not show a specific trend with oligomer loadings (Figures S10b and S11b). The AUPs of inter β-strand signals were calculated after the knowledge-based spectrum decomposition into three signals corresponding to tissue, inter β-sheet and inter β-strand. The decomposed signals matched quite well with their corresponding quantity- and region-matched reference standards (Figure 5d). The AUPs of inter β-strand signals, decomposed from composite spectra associated to olfactory bulb and medial temporal lobe, correlated well with their respective ground truths with the Pearson correlation coefficient above 0.93 (Figure 5f).

The study confirms that the presence of even 50 µg of oligomer results in a quantitatively measurable change in X-ray signal, even with a laboratory source. Irrespective of the type of aggregates (fibril or oligomer), the variation in the aggregate burden in the tissue changes the X-ray spectrum amplitude. The impact of aggregates on X-ray spectrum width is not systematic, suggesting that spectrum width may not serve as a quantitative metric to track the incremental deposition of aggregates. The accuracy of the material decomposition approach, based on knowledge-based peak deconvolution and quantification, is confirmed by the similarity between decomposed signals and their corresponding ground truths, along with the high Pearson’s correlation coefficient.

### 3.5. XβAI for comparing regional aggregate burden between tissue regions

Next, we identified ten distinct regions of interest (ROIs) within sheep brain tissue and examined their X-ray scattering profiles both with and without a constant oligomer burden. The investigated brain tissue regions spanned from the brain stem to the medial temporal lobe, cortex, and cerebellum, as indicated in Figure 6a. All ten tissue regions scattered the X-ray beam, and the resultant X-ray spectra were centered at around *q*_*c,tissue*_ ∼12.7 ± 1.5 *nm*^−1^ (Figure 6b). The tissue scattering volume (*V*_*tissue*_), calculated by assuming a cylindrical shape for the ROI X-ray interacting volume with the beam size as the radius and tissue thickness as the height, exhibited variation across different brain regions. Among the ROIs studied, the olfactory bulb exhibited the smallest thickness, while the occipital cortex was identified as the thickest tissue. The spectrum amplitude increased with tissue thickness and the spectrum width did not show any specific correlation with the thickness (Figure S12). The *V*_*tissue*_ of different tissue regions and their X-ray spectrum area calculated by integrating each spectrum between *q* ranges from 4.2 *nm*^−1^ to 25 *nm*^−1^ exhibited similar trend (Figure 6c). This suggests that the structure of X-ray scatters present in all ROIs is qualitatively comparable, but there is heterogeneity in the number of scatters across the tissue regions.

**Figure 6:**
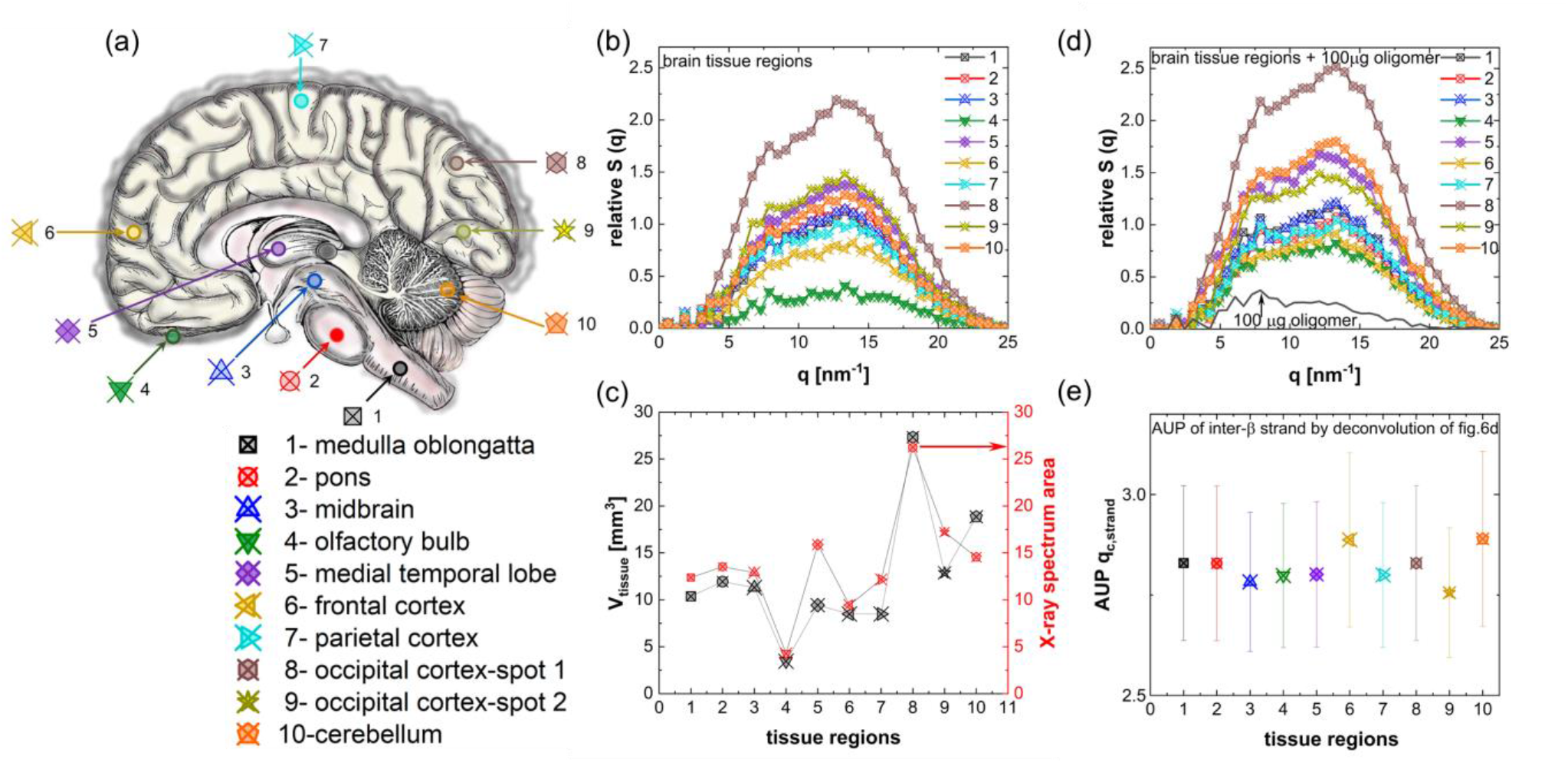
The wide-angle X-ray scattering properties of sheep brain tissues from various brain regions were investigated, both with and without the oligomer burden. (a) The schematic illustrates the brain tissue regions subjected to X-ray scanning. (b) The WAXS profiles of tissues from distinct brain regions are presented. The regions are labeled numerically from 1 to 10, and the corresponding regions can be identified in Figure 6a. (c) The correlation between tissue scattering volume (V_tissue_), calculated by assuming a cylindrical shape for the ROI X-ray interacting volume, with the beam size as the radius and tissue thickness as the height, and the total area under the WAXS spectrum for each tissue ROI is illustrated. (d) The WAXS profiles following the introduction of 100 µg BSA oligomer to the ten different ROIs. The Y-axis scale is maintained same for Figures 6b and 6d, facilitating the straightforward interpretation of the impact of a 100 µg oligomer burden on all 10 tissue ROIs. (e) The area under the peak (AUP) estimated for all ten tissue ROIs with 100 µg oligomer, by integrating the inter-β strand peak (q_c,strand_ ∼13.3 nm^−1^) decomposed from composite spectra. The AUPq_c,strand_ ∼13.3 nm^−1^ are quantitatively similar for all ten ROIs irrespective of differences in the composite spectra. The relative errors are estimated based on the standard of reference value calculated from inter-β strand peak of 100 µg oligomer alone. The area under the peak (AUP) calculated for all ten tissue ROIs with 100 µg oligomer, by integrating the inter β-strand peak (q_c,strand_∼13.3 nm^−1^) decomposed from composite spectra. The 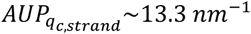 is quantitatively similar for all ten ROIs, regardless of differences in the composite spectra. Relative errors are estimated based on the standard reference value calculated from the inter β-strand peak of 100 µg oligomer alone.

#### 3.5.1. Fixed oligomer loading at different tissue regions

To assess the capability of X-ray scattering technique in detecting protein aggregates from various brain tissue regions and to estimate the accuracy in decomposing aggregate scattering profiles from diseased tissues, we introduced 100 μg oligomer into tissues from ten different ROIs. Addition of oligomer increased the spectrum amplitude compared to their corresponding tissue alone spectra (compare Figures 6b and 6d). The influence of oligomer burden on spectrum amplitude was notably pronounced in low-scattering volume tissues, such as the olfactory bulb. The X-ray profiles of tissue, inter-β strand, and inter-β sheet were deconvolved from the composite spectrum based on peak location knowledge as described before. The decomposed inter-β strand and inter-β sheet signals from ten different composite spectra were both qualitatively and quantitatively similar, showing comparable signals with respect to the X-ray spectrum of 100 µg oligomer alone. The decomposed tissue signals were different for each ROIs due to difference in *V*_*tissue*_. Figure 6e concludes that the area under the peak of the decomposed inter-β strand peaks (*q*_*c,strand*_ ∼13.3 *nm*^−1^) were quantitatively similar for all ten ROIs, irrespective of differences in the composite spectra. The relative errors, calculated based on the ground truth value, were within 20%. Therefore, we successfully quantified the same amount of oligomer loading from different tissue surroundings with varying *V*_*tissue*_. The findings were further validated by performing a similar study with 6 mg oligomer loading embedded in ten different tissue regions (Figure S13 in supporting information). The observations were qualitatively similar to those observed in tissues with 100 µg oligomer. Additionally, the effect of aggregate burden on tissue scattering profiles was more prominent in terms of spectrum amplitude with an increase in aggregate burden (compare figures 6d and S12a). This study concludes that the material decomposition approach can accurately recover protein aggregate signals from heterogeneous brain tissue regions.

#### 3.5.2. Quantification of regional aggregate density in different tissue regions

To quantify the regional accumulation of aggregates, we propose a quantitative metric, X-ray cross-β aggregate index (XβAI). The *XβAI*_*r*_ is the ratio of area under the peak (AUP) of inter β-strand peak to the AUP of tissue signal deconvolved from raw composite spectrum (equation 10). The index calculates the aggregate burden per tissue scattering volume and it represents the aggregate density within the volume of region of interest. For instance, *XβAI*_*r*_ was calculated for ten different tissue regions with 100 µg and 6000 µg oligomer amounts (Figure 7a). For a fixed oligomer amount, the regional aggregate density varied corresponding to the tissue volume (*V*_*tissue*_).

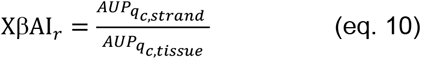

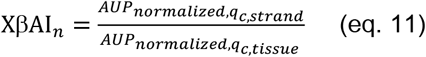

The index, *XβAI*_*n*_ was used to represent the change in the aggregate amount within a ROI volume. The *XβAI*_*n*_ is the ratio of AUP of inter β-strand peak to the AUP of tissue signal deconvolved from normalized composite spectrum (equation 11). The composite spectrum was normalized with the photon count at *q*_*c,tissue*_ of deconvolved tissue signal in the raw composite spectrum. Normalization with tissue signal homogenized the differences between tissue signals originated from the differences in *V*_*tissue*_. For example, in Figure 7b, the differences between normalized spectra of olfactory bulb tissue with oligomers are entirely arising from the differences in the oligomer burden and the AUPs of tissue signals are same for all compositions. The *XβAI*_*n*_ was zero for ‘healthy’ tissue with no oligomers and the index increased with increase in oligomer amount from 50 µg to 6000 µg. Our data suggests that the index, *XβAI*_*n*_ could serve as a potential quantitative metric for monitoring aggregate burden in longitudinal studies and for comparing tissues across different brain regions, providing a standardized measure by normalizing the tissue signals. Furthermore, the material decomposition approach effectively separates individual component signals in the composite spectrum. As a result, the proposed methodology eliminates the need for subtracting healthy tissue signals as background to isolate aggregate-alone signals. The prior reported protein aggregate X-ray scattering signals from ex-vivo brain tissue of various NDDs were obtained by subtracting the scattering signals of ROI matched healthy tissues. However, it is impossible to accurately match *V*_*tissue*_ for each ROIs and that can lead to left over tissue signals in the aggregate signals especially in the *q* regions correspond to the inter-β strand signals of aggregates. The previously reported X-ray scattering signals of protein aggregates from ex-vivo brain tissue in various neurodegenerative diseases were derived by subtracting the scattering signals of ROI-matched healthy tissues. However, achieving accurate matching of *V*_*tissue*_ for each ROI is challenging, potentially resulting in residual tissue signals within the aggregate signals, particularly in the *q* regions corresponding to the inter-β strand signals of aggregates. Consequently, this mismatch may result in an overestimation of the aggregate burden. The suggested material decomposition method, without the need for defining reference regions, enhances the reliability of estimating aggregate burden using wide-angle X-ray scattering techniques.

**Figure 7:**
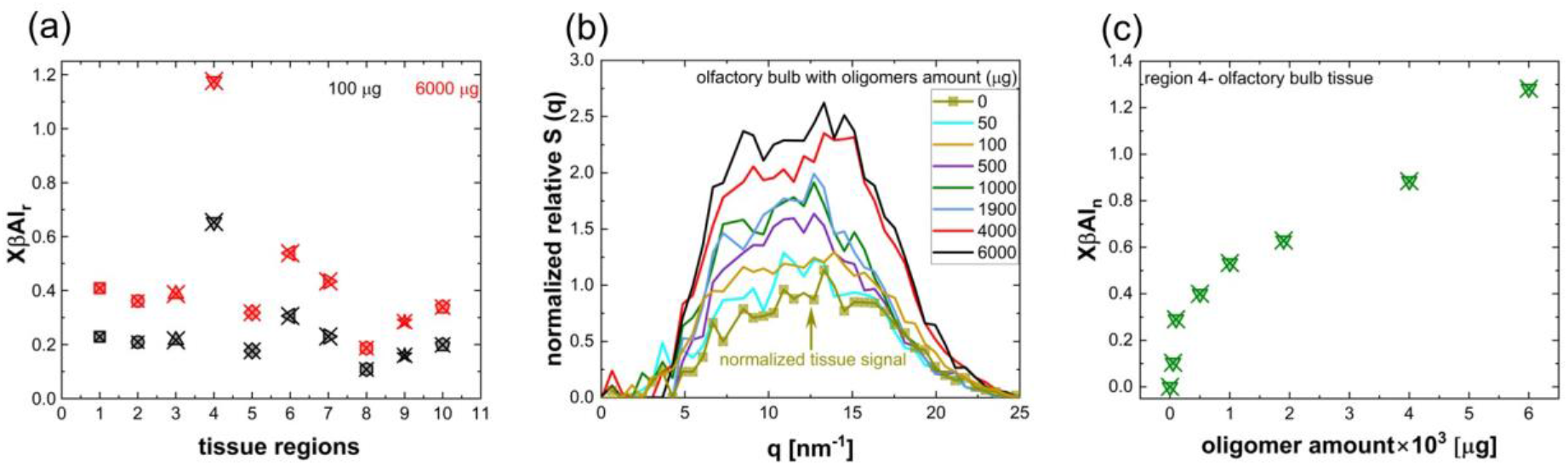
The quantification of regional aggregate burden in tissues, expressed as the X-ray Cross-β Aggregate Index (XβAI), is calculated both with (XβAI_n_) and without (XβAI_r_) spectrum normalization using the decomposed tissue signal. (a) The XβAI_r_, estimated for ten different tissue ROIs with oligomer burden of 100 µg and 6 mg. (b) The normalized spectra of olfactory bulb tissue, both with and without varying oligomer burden in the same ROI, are shown. The spectra were normalized based on the maximum amplitude of the tissue signal decomposed from the raw composite spectrum. (c) The XβAI_n_ estimated for olfactory bulb with varying oligomer burden.

## 4. Discussion

The present wide-angle X-ray scattering (WAXS) studies focusing on the quantification of oligomers and fibrils in isolation and within diverse brain tissues, validate the effectiveness of the X-ray scattering technique employing laboratory-source X-rays. This method successfully identified cross-β signals emerging from oligomer and fibrillar models ranging from 50 µg to 6 mg. This was achieved even in oligomer models containing 18% β-sheet (Figures 4 and 5). The previously reported aggregate quantities in human brain tissues from distinct neurodegenerative diseases exhibit a broad spectrum, spanning from nanograms to milligrams. This variance is contingent upon the type of aggregate, disease staging, specific brain region and tissue volume, as documented in various studies [40–44]. For instance, Anagnostou et al. reported approximately 7 (± 3.03) and 25 (± 30.21) nanograms of aggregated α-synuclein per milligram of tissue in the caudate nucleus and putamen regions, respectively, in patients with Parkinson’s disease with an average disease duration of 5 years [40]. Contrarily, age and sex-matched tissues devoid of neurodegenerative disease exhibited α-syn aggregate levels of 1.5 (± 0.15) and 3.4 (± 1.34) ng/mg tissue in the caudate nucleus and putamen, respectively. In another study by Roberts et al., who fractionated frontal cortical grey matter from the postmortem brains of 78 (±9) year-olds with Alzheimer’s disease (AD), an average of 6.5 (±5) mg total amyloid-β was reported, as opposed to 1.7 (±2.3) mg in control brains [41]. The total Aβ concentration correlated with a 5 µg/g of grey matter for the PET SUVr threshold for Aβ positivity (SUVr=1.4) and 11.2 µg/g of grey matter for mean Alzheimer’s disease dementia SUVr values of 2.3. The Aβ aggregates accumulation rate reported was 28 ng/h. In our study, the signal-to-noise ratio (SNR) estimation, based on Equation 4 for a 50-µg oligomer was around one (Figure S5 in supporting information), with values increasing with aggregate quantity. Further enhancements in SNR are achievable through optimization of experimental conditions including optimization of X-ray exposure conditions, enlarging the scattering volume of the region of interest, primarily by adjusting X-ray beam dimensions, and by fine-tuning detector photon detection thresholds and wavevector resolutions. Additionally, the development of X-ray scattering contrast agents including nanoparticulate can also further increase the detectability of very low/sparse aggregates [45].

The X-ray scattering profiles of oligomers and fibrils, when present in clinically relevant quantities, exhibit overlaps between inter-β strand and inter-β sheet peaks. This stands in contrast to the typical amyloid peaks characterized by well-defined maxima at peak locations *q*_*c,sheet*_ = 6.28 *nm*^−1^ and *q*_*c,strand*_ = 13.36 *nm*^−1^ (Figure 4). The polymorphism observed in the spectra primarily manifests as variations in the ratios of peak widths and peak areas between inter-β strand and inter-β sheet (Table S1). Notably, existing literature reports differences in X-ray diffraction cross-β signals both within pathogenic strains of a patient and between different patients [28,29]. For instance, Araki et al. demonstrated distinctions in the sharpness of X-ray diffraction spectra of Lewy bodies in midbrain tissue identified through antibody staining to α-synuclein from a Parkinson’s patient and among age and sex-matched patients [29]. Their findings encompassed diffraction spectra featuring sharp inter-β strand and sheet peaks, as well as broader spectra lacking well-defined maxima for inter-β sheet peaks of α-synuclein in tissue. Similarly, Liu et al. provided evidence of structural polymorphisms in amyloid fibrils within and among plaques of an individual with AD without dementia [28]. They identified cross-β signals of Aβ fibrils with inter-β strand distance sharply peaked at 0.47 nm (*q*_*c,strand*_ = 13.36 *nm*^−1^) or as a doublet with varying intensity ratios at 0.465 nm (*q*_*c,strand*_ = 13.51 *nm*^−1^), and 0.475 nm (*q*_*c,strand*_ = 13.23 *nm*^−1^). Their study concluded that peak area ratios are linked to plaque packing densities, with the highest peak area ratio of 0.475/0.465 were found near plaque cores. This ratio decreased while scanning externally to the plaque core and for diffuse plaques. In a prior study, we also reported alterations in the ratios of inter-β strand to inter-β sheet peak areas corresponding to changes in aggregate packing density [32].

The polymorphic nature of biomarker aggregates in neurodegenerative diseases (NDDs) extends beyond the realm of X-ray cross-β signals, encompassing distinctions observed through microscopy imaging and seed aggregation assays. Cryo-electron microscopy has revealed structural disparities between α-synuclein aggregates in multiple system atrophy (MSA) and dementia with Lewy bodies (DLB) [46,47]. Furthermore, Martinez Valbuena et al. have reported substantial heterogeneity in the seeding activity of MSA-derived α-synuclein across distinct brain regions within the same individual and attributed these variations to potential structural and biochemical heterogeneity of α-synuclein [48].

Remarkably, our analysis of previously reported cross-β signals of Aβ fibrils, oligomers, paired helical filaments of tau, and Lewy bodies in brain tissues from NDD patients or from animal models, as well as their in-vitro recombinants, reveals that structural polymorphism minimally impacts the peak locations of inter-β strand and inter-β sheets (Table S1). The averaged peak locations derived from various cross-β signals were *q*_*c,sheet*_ = 6.03 ± 0.64 *nm*^−1^ and *q*_*c,strand*_ = 13.36 ± 0.57 *nm*^−1^. In the majority of cases, inter-β strand peaks were consistently centered around 13.3 *nm*^−1^ with an inter-β strand spacing of 0.47 nm. We believe the deviations to *q*_*c,strand*_ = 15 or 12 *nm*^−1^ can also stem from residual tissue scattering signals due to inadequate background subtraction during composite spectrum pre-processing. Structural heterogeneities have a discernible impact on the area and width ratios of peaks corresponding to β-sheets and strand spacings (as outlined in Table S1). The spectrum heterogeneity may arise from diverse factors such as variations in clinical histories, maturity stages of deposits, coexistence of different aggregation levels, aggregate packing density, β-sheet packing order, structural organization within the surrounding tissue, and variation in fibril twist. Consequently, the decomposition of aggregate X-ray signals from composite spectra based on peak location knowledge, along with the assignment of peak area and width as floating variables are justified.

The brain tissue phantom study provides compelling validation that the area under the peak (AUP) of the inter-β strand effectively represents the percentage of aggregates within the tissue-aggregate composite (Figure 2). The existing literature suggests the absence of a peak centered on the inter-β strand spacing in healthy brain tissues. For example, Liu et al., reported the absence of a peak corresponding to a 0.47 nm spacing in 2500 X-ray diffraction patterns collected by spatial X-ray diffraction mapping of healthy postmortem brain tissue without known NDDs [28]. Moreover, previous studies have identified amyloid, and Lewy body-positive cases based on the presence of diffraction reflections at 0.47 nm in postmortem human brain tissues (Table S1). The peak corresponding to inter-β sheet spacing (∼1 nm) was either absent or negligibly small in synchrotron-based X-ray diffraction signals reported thus far (Table S1). Therefore, it is difficult to establish a correlation between AUP of inter-β sheet spacing against the aggregate burden. In our phantom study, the inter-β strand peak’s AUP fraction with respect to the total AUP of the phantom signal exhibited a linear correlation with the aggregate fraction in the phantom matrix (*f*_*agg,true*_), as illustrated in Figure 2d.

The sheep brain tissues displayed distinct X-ray scattering patterns and the intensity of X-ray scattering displayed a nearly linear increase with varying tissue thickness. The independence of spectrum width on tissue thickness suggests a uniform distribution of similarly structured scatters throughout different brain regions. However, significant variability in the number of such scatters across different tissue regions is observed. The spectra maxima, approximately around *q*_*c,tissue*_ ∼12.6 *nm*^−1^ suggest that primary scatters within the tissues may be attributed to lipids. We anticipate similar scattering properties for human brain tissues due to high degree of homology between the human and sheep brains, including their convoluted cortices. The improved animal-to-human translation is demonstrated using higher-order mammals with more complex and comparable neuroanatomy compared to commonly used small laboratory animals [49,50]. Historically, the sheep brain has served as an animal model for PD [51,52], to study the formation of Alzheimer’s-related neurofibrillary tangles [53,54] and amyloid plaques [55] and as a model system for Huntington’s disease [56]. Furthermore, amino acid sequences of Alzheimer’s disease (AD)-associated proteins, such as amyloid precursor protein (APP), and fragments of APP found in aged sheep, have been reported to mirror those commonly identified in humans [55].

The WAXS signals from clinically relevant quantities of aggregates had low amplitude (or low echo) compared to the tissue spectrum in majority of synthetically diseased tissues from different brain regions due to large quantities of tissue scatters within the ROI scattering volume. This holds true for both oligomer and fibrillar model aggregates studied here. However, the scattering volume of olfactory bulb tissue was lower than tissues from all other brain regions studied here. Therefore, the effect of oligomers on olfactory bulb scattering profile was more easily detectable than any other tissue regions. The addition of aggregate to the tissue increased the spectrum amplitude and an initial increase in the width to the low *q* side of the spectrum (Figures 6, S7-S13). For example, the addition of 100 µg fibril to the tissue from frontal cortex increased the spectrum width to the left side of the spectrum half maximum (between *q* = 2.4 *to* 12.6 *nm*^−1^) by 20% (Figure S7). The spectrum broadening to low *q* side arises from the presence of peak corresponding to the inter-β sheet spacing. In addition to that the peak corresponds to inter-β strand spacing increases the spectrum amplitude with increase in aggregate burden.

The signal decomposition by Gaussian mixture model with prior knowledge of X-ray scattering peak locations of tissue, water, inter-β sheet and inter-β strand spacings of aggregates successfully recovered the low echo aggregate signals which were merged inside the composite spectrum. The effectiveness of the proposed material decomposition method in detecting and quantifying the different nanoscopic X-ray scattering structures in a material from their scattering spectrum was tested under various realizations. The method successfully decomposed the X-ray spectrum from (i) human brain grey and white matter tissues (Figure 2) (ii) brain tissue phantoms with 6 different tissue-aggregate proportions (Figure 3) (iii) 7 different quantities (50 µg to 6 mg) of model oligomer (Figures 4a,b) (iv) 6 different quantities of model fibril (100 µg to 6 mg, Figures 4c,d) (v) sheep brain tissues from 10 different brain regions varying from brain stem-to-cortex-to-cerebellum (Figure 6a,b) (vi) frontal cortex tissue with 7 different quantities of fibrillar model (0 µg to 6 mg, Figures 5a,b,e) (vii) olfactory bulb tissue with 8 different quantities of oligomer model (0 µg to 6 mg, Figures 5c,d,f) (viii) medial temporal lobe tissue with 6 different quantities of oligomer model (0 µg to 1.9 mg, Figures 5f, S9) (ix) 100 µg oligomer variation in tissue from 10 different brain regions (Figures 6d,e) (x) 6 mg oligomer variation in tissue from 10 different brain regions (Figure S13). The results demonstrated the effectiveness of the proposed method under all realizations described above. Additionally, the method was successful for both synchrotron and laboratory-source based X-ray scattering spectrum. The quantification of nanostructures in terms of the AUP of corresponding decomposed signal correlated very well with their standard of reference with Pearson’s correlation coefficient (*r*) above 0.9. In our study the ground truths were established from literature reports or from independent studies on each component of a material after quantitatively and qualitatively matching. The relative error for each realization estimated based on ground truths were mostly within 20%. The relative error between mean values was contributed by different factors including differences in nanostructure packing with respect to the surrounding along with standard deviation between repetitive measurements, minor deviations in sample preparation and X-ray scanning locations between ground truths and composite samples, signal noise and model fitting errors.

Moreover, the aggregate signal recovery from composite spectrum by decomposition method is independent of tissue heterogeneity. The AUPs of decomposed inter-β strand signals were similar for a fixed quantity aggregate collected from any part of the brain region and the values were comparable with their ground truths (Figures 6 and S13). Excitingly, this suggests the possibility of regional detection and accurate quantification of biomarker aggregates in the tissues from any part of the brain regions using WAXS. Prior studies have shown distinct biomarker distribution pattern in the central nervous system for different NDDs [57–59]. For example, studies have shown olfactory dysfunction is observed prior to the motor symptoms in PD [60]. This suggests that the PD pathology may begin in the olfactory bulb. Aggregation of α-synuclein is observed in the olfactory centers such as the olfactory bulb (OB) and the anterior olfactory nucleus during the early stages of PD [61]. This observation also underscores the importance of investigating how abnormal α-synuclein affects the function of the OB and olfaction may provide crucial information for the pathological processes in PD. Recent in vivo quantitative neuroimaging investigations have shown brain Aβ accumulation to occur initially in cerebral regions and spread from neocortex to allocortex to brainstem, eventually reaching the cerebellum [58,62]. The neurofibrillary tau tangles (NFTs) are more numerous in medial temporal lobe regions associated to memory functions. The NFTs in AD patients are reported to be initially limited to the hippocampus, entorhinal cortex and adjacent limbic structures. Later NFTs spread to temporal neocortex-to-lateral temporal cortex-and finally to frontal or parietal neocortex [63,64]. Figure 5c and Figure S9 suggest that the oligomers in olfactory bulb and medial temporal lobe can be detected by WAXS. Additionally, the regional increase in oligomer burden was accurately quantified by material decomposition. Similar observation was found for fibrils embedded in frontal cortex tissue (Figure 5a,b).

Importantly, we proposed a new metric to quantify the biomarker burden in NDDs based on the X-ray scattering signal of cross-β structure of protein aggregates. Several X-ray microdiffraction studies have detected the cross-β signals of amyloid-β plaques, neurofibrillary tangles, and Lewy body deposits in ex-vivo human brain tissues from various NDD patients including Alzheimer’s disease, Parkinson’s disease, Down syndrome, frontotemporal lobar degeneration, and multiple system atrophy (Table S1). However, so far, the X-ray diffraction method was used for the qualitative analysis of postmortem brain tissues based on the presence or absence of the inter-β strand peak in the one-(1-D) or two-dimensional(2-D) diffraction data. Our results suggest the possibilities of quantitative analysis along with the regional or global estimation of pathogenic aggregates using wide angle X-ray scattering technique. The aggregate burden can be quantified as the area under the peak of inter-β strand peak. However, the AUP values are dependent on the measurement conditions and cannot compare across different experimental conditions. Therefore, to harmonize the aggregate quantities measured under varying measurement conditions, we proposed the cross-β aggregate index (XβAI_*r*_) by taking the ratio of AUPs of decomposed inter-β strand signal to the AUP of decomposed tissue signals from the same composite spectrum. The index calculated after the normalization of composite spectrum with the decomposed tissue signal characteristic eliminates the variations on a tissue over the period (XβAI_*n*_). Therefore, the XβAI_*n*_ is independent of ROI definition suggests that the aggregate burden could be directly comparable between different brain regions. Additionally, the percentage aggregate burden in a scattering volume can also be estimated using equation 12:

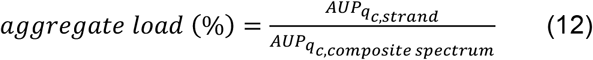

The % aggregate load represents the total percentage of inter-β strand spacing present within the composite spectrum of a ROI arising from the scattering of tissue, inter-β strand- and inter-β sheet-spacings of X-ray scattered aggregates. The estimation of XβAI can be for the specific brain region set by the scattering volume as well as for the global estimate by combining composite spectra acquired from different ROIs.

The major advantages of XβAI over the previously discussed quantitative metrics based on PET, as well as fluid biomarkers include the following: the proposed methodology does not require the definition of a reference region, there is no need for spatial registration on MRI, and it does not require a contrast agent to obtain the X-ray scattering signals of aggregates. Furthermore, it offers the potential for regional as well as global biomarker estimation. The proposed methodology can be used as a virtual histology (VH) tool for the detection and quantification of aggregates in postmortem brain tissues. The X-ray absorption-based 3D virtual histology for bone, dental studies, heart, and kidney were reported previously [65], and it would be interesting to study the X-ray scattering based VH for amyloidoses. Compared to the immunohistochemistry methods, the WAXS study will allow the aggregate estimation from thicker tissue samples (several millimeter) than the immunostained microtomed (<50 µm) samples. The mapping of aggregates is expected to be faster than the histology due to non-requirement of sample preparations including segmentation and staining prior to the experiments. Additionally, the XβAI can be estimated using synchrotron as well as laboratory X-ray sources. The synchrotron-based measurements can further reduce the total measurement time compared to the lab source X-ray.

Recent reports indicate the potential of the spectral X-ray scattering technique to capture target signals from objects as thick as the human head without the need for a radiotracer or any other contrast agents [66]. Initial phantom and animal model studies, along with the simulation results, suggest that the combination of a photon counting detector and a polychromatic X-ray source, with the potential to tune the X-ray energy range, permits to probe the scattering of targets from thicker objects in contrast to conventional X-ray scattering technique. Dahal et al. demonstrated a strong correlation between the estimated brain amyloid burden in Alzheimer’s disease (AD) mice models using the spectral X-ray scattering technique and histological results [35]. Following the success of the X-ray scattering based technique in clinical studies in the future, XβAI_*n*_ can be a potential metric for longitudinal studies to monitor the deposition of protein aggregates.

The X-ray scattering technique, owing to the generalized cross-β structure of hallmark biomarkers such as amyloid-β, α-synuclein, and tau in Alzheimer’s disease, Parkinson’s disease, and tauopathies, respectively, emerges as a versatile method for the detection and quantification of multimodal biomarkers. The X-ray signal of inter-β strand spacing, in conjunction with the clinical representation of regional biomarker distribution in the brain and associated clinical symptoms or functional assessments, holds the potential to discriminate between different neurodegenerative diseases in the context of cross-β structure-based biomarker detection and quantification methods. Discrimination of NDDs based on the regional distribution of biomarkers has been implied previously as well. For instance, the increased α-synuclein targeting PET tracer uptake is reported in the basal ganglia region for Multiple system atrophy-parkinsonian type (MSA-P) compared to normal and PD cases [67]. In contrast, Multiple system atrophy - cerebellar subtype (MSA-C) cases showed tracer retention in the cerebellar white matter and cerebellar peduncles. Regional structural differences between AD, PD and FTD are also reported previously. For example, in DLB, the hippocampus is typically preserved, in contrast to severe hippocampal atrophy in AD and temporal pole atrophy in FTD [68]. Moreover, the deposition of protein aggregates is not exclusive to neurodegenerative diseases (NDDs). Such protein aggregation has been reported in other amyloidoses affecting organs such as the liver, kidney, spleen, and heart. The generic cross-β substructure is reported for aggregate deposits of non-NDDs including transthyretin (TTR) amyloidosis [69] and serum amyloid A rheumatoid arthritis [70]. Indeed, tracking TTR protein deposits using the proposed X-ray scattering method could be particularly intriguing, especially considering recent FDA approval of Wainua™ for the treatment of polyneuropathy associated with hereditary transthyretin-mediated amyloidosis in adults [71]. Consequently, the proposed X-ray scattering-based material decomposition and aggregate quantification method can be expanded for in vivo biomarker imaging studies beyond NDDs.

The proposed X-ray scattering-based material decomposition method can also be extended to study other tissue alterations, such as those occurring in diabetes mellitus due to biochemical reactions, and in traumatic brain injury by tracking changes in myelin nanostructure. The current study is limited to brain tissue phantoms and sheep brain tissues. However, further studies on human brain tissues with protein aggregates linked to different diseases and disease severities will be needed to confirm its effectiveness in the detection, differentiation, and quantification of diverse protein aggregates. In the future, it would also be interesting to establish XβAI cut-off values for different disease stages and their accuracy estimates with respect to regulatory-accepted reference standards.

## Conclusions

In summary, our study introduces a novel quantitative metric based on X-ray scattering for the comprehensive assessment of protein aggregates associated with diverse amyloidoses, including neurodegenerative diseases. Unlike current imaging modalities used for amyloidoses, the X-ray scattering technique specifically targets the inherent nanoscopic cross-β structure of protein aggregates, eliminating the need for a radiotracer or other contrast agents. Our findings demonstrate the potential of the wide-angle X-ray scattering technique to detect and quantify both oligomeric and fibrillar aggregates across various brain regions, suggesting its utility as an early-stage disease detection method. The correlation between the area under the peak of inter-β-strand signals and aggregate burden was robust. The proposed material decomposition approach effectively separates convoluted signals in the composite spectrum, enabling the recovery of aggregate signals from intricate tissue scattering profiles without the necessity of defining a reference region. Validation under various realizations confirmed the method’s accuracy, with decomposed inter-β-strand signals exhibiting strong correlation with respect to the ground truth (Pearson correlation coefficient consistently above 0.9). Consequently, the proposed X-ray Cross-β aggregate index stands as a promising tool for quantifying regional protein aggregate burden and understanding their deposition pathways.

## List of abbreviations

AD: Alzheimer’s disease
AUP: Area under the peak
Aβ-42: Amyloid β−42
Aβ: Amyloid β
BSA: Bovine serum albumin
CdTe: Cadmium telluride
CERAD: Consortium to establish a registry for Alzheimer’s disease
CI: Confidence interval
CL: Centiloid
CSF: Cerebrospinal fluid
CT: Computed tomography
CU: Cognitively unimpaired
FDA: U.S. Food and Drug Administration
FTD: Frontotemporal lobar degeneration
FWHM: Full width half maximum
FWTM: Full width tenth maximum
GCI: Glial cytoplasmic inclusions
LB: Lewy bodies
LGAB: β-lactoglobulin genetic A and B mixture
MCI: Mild cognitive impairment
MRI: Magnetic resonance imaging
MSA: Multiple system atrophy
NDDs: neurodegenerative diseases
NFT: Neurofibrillary tangles
OB: Olfactory bulb
PD: Parkinson’s disease
PET: Positron emission tomography
PS: Parkinsonian syndromes
ROI: Region of interest
SDD: Sample to detector distance
SNR: Signal-to-noise ratio
SPECT: Single photon emission computed tomography
SUVr: Standardized uptake value ratio
sXS: Spectral X-ray scattering
TTR: Transthyretin
VH: Virtual histology
WAXS: Wide-angle X-ray scattering
XβAI: X-ray cross-β amyloid index

## Declarations

### Ethical approval and consent to participate

Not applicable.

### Consent for publication

Not applicable.

## Acknowledgements

KS acknowledges funding by appointments to the Research Participation Program at the Center for Devices and Radiological Health and the Center for Drug Evaluation and Research administered by the Oak Ridge Institute for Science and Education through an interagency agreement between the U.S. Department of Energy and the U.S. Food and Drug Administration. KS acknowledges John Dennis from FDA-Center for Biologics Evaluation and Research, David Rotstein from FDA-Center for Veterinary Medicine and Ilyas Saytashev from FDA-Center for Devices and Radiological Health for the sheep brain anatomy discussion. KS acknowledges Bahaa Ghammraoui from FDA-Center for Devices and Radiological Health for providing spectral X-ray lab facility and for discussion on performance metrics for material decomposition. The mention of commercial products herein is not to be construed as either an actual or implied endorsement of such products by the Department of Health and Human Services. This is a contribution of the Food and Drug Administration and is not subject to copyright.

## Author contributions

KS: conceptualization, methodology, investigation, data curation, writing-original draft preparation

ED: conceptualization, funding acquisition, supervision on spectral X-ray scattering experiments, writing-review & editing.

AB: conceptualization, funding acquisition, overall supervision, writing-review & editing, approval of final version of manuscript

## Availability of data and materials

All data supporting the conclusions of this article are included within the article and in supporting information provided.

## REFERENCES

[1] https://www.accessdata.fda.gov/drugsatfda_docs/label/2022/022454s010lbl.pdf

[2] https://www.accessdata.fda.gov/drugsatfda_docs/label/2019/200655s000lbl.pdf

[3] https://www.accessdata.fda.gov/drugsatfda_docs/label/2012/202008s000lbl.pdf

[4] https://www.accessdata.fda.gov/drugsatfda_docs/label/2014/204677s000lbl.pdf

[5] https://www.accessdata.fda.gov/drugsatfda_docs/label/2017/203137s008lbl.pdf

[6] Pemberton HG, Collij LE, Heeman F, Bollack A, Shekari M, Salvadó G, et al. Quantification of amyloid PET for future clinical use: a state-of-the-art review. Eur J Nucl Med Mol Imaging 2022;49:3508–28. 10.1007/s00259-022-05784-y.

[7] https://www.accessdata.fda.gov/drugsatfda_docs/summary_review/2023/761269Orig1s000SumR.pdf

[8] https://www.accessdata.fda.gov/drugsatfda_docs/nda/2021/761178Orig1s000MedR_Redacted.pdf

[9] Villemagne VL, Leuzy A, Bohorquez SS, Bullich S, Shimada H, Rowe CC, et al. CenTauR: Towards a Universal Scale and Masks for Standardizing Tau Imaging Studies. Neurology; 2023. 10.1101/2023.03.22.23287009.

[10] Heeman F, Yaqub M, Lopes Alves I, Heurling K, Bullich S, Gispert JD, et al. Simulating the effect of cerebral blood flow changes on regional quantification of [18F] flutemetamol and [18F] florbetaben studies. J Cereb Blood Flow Metab 2021;41:579–89. 10.1177/0271678X20918029.

[11] Amadoru S, Doré V, McLean CA, Hinton F, Shepherd CE, Halliday GM, et al. Comparison of amyloid PET measured in Centiloid units with neuropathological findings in Alzheimer’s disease. Alz Res Therapy 2020;12:22. 10.1186/s13195-020-00587-5.

[12] Doré V, Bullich S, Rowe CC, Bourgeat P, Konate S, Sabri O, et al. Comparison of [18F]‐florbetaben quantification results using the standard Centiloid, MR‐based, and MR‐less CapAIBL® approaches: Validation against histopathology. Alzheimer’s &amp; Dementia 2019;15:807–16. 10.1016/j.jalz.2019.02.005.

[13] for the Alzheimer’s Disease Neuroimaging Initiative, for the ALFA Study, Salvadó G, Molinuevo JL, Brugulat-Serrat A, Falcon C, Grau-Rivera O, et al. Centiloid cut-off values for optimal agreement between PET and CSF core AD biomarkers. Alz Res Therapy 2019;11:27. 10.1186/s13195-019-0478-z.

[14] Whittington A, Gunn RN. Amyloid Load: A More Sensitive Biomarker for Amyloid Imaging. J Nucl Med 2019;60:536–40. 10.2967/jnumed.118.210518.

[15] Leuzy A, Lilja J, Buckley CJ, Ossenkoppele R, Palmqvist S, Battle M, et al. Derivation and utility of an Aβ-PET pathology accumulation index to estimate Aβ load. Neurology 2020;95:e2834–44. 10.1212/WNL.0000000000011031.

[16] Pegueroles J, Montal V, Bejanin A, Vilaplana E, Aranha M, Santos‐Santos MA, et al. AMYQ: An index to standardize quantitative amyloid load across PET tracers. Alzheimer’s &amp; Dementia 2021;17:1499–508. 10.1002/alz.12317.

[17] https://www.accessdata.fda.gov/drugsatfda_docs/label/2020/212123s000lbl.pdf

[18] https://www.accessdata.fda.gov/cdrh_docs/reviews/K231348.pdf

[19] https://www.accessdata.fda.gov/cdrh_docs/reviews/K221842.pdf

[20] https://www.accessdata.fda.gov/cdrh_docs/reviews/DEN200072.pdf

[21] https://alz.org/media/Documents/scientific-conferences/Clinical-Criteria-for-Staging-and-Diagnosis-for-Public-Comment-Draft-2.pdf?_gl=1*jirfag*_ga*OTk0NDExMDAzLjE2NTE2ODAxNTY.*_ga_QSFTKCEH7C*MTcwMzkxMDMwOC44NC4wLjE3MDM5MTAzMDguNjAuMC4w*_ga_9JTEWVX24V*MTcwMzkxMDMwOC44NC4wLjE3MDM5MTAzMDguNjAuMC4w

[22] Cullen NC, Janelidze S, Mattsson‐Carlgren N, Palmqvist S, Bittner T, Suridjan I, et al. Test‐retest variability of plasma biomarkers in Alzheimer’s disease and its effects on clinical prediction models. Alzheimer’s &amp; Dementia 2023;19:797–806. 10.1002/alz.12706.

[23] Thijssen EH, La Joie R, Strom A, Fonseca C, Iaccarino L, Wolf A, et al. Plasma phosphorylated tau 217 and phosphorylated tau 181 as biomarkers in Alzheimer’s disease and frontotemporal lobar degeneration: a retrospective diagnostic performance study. The Lancet Neurology 2021;20:739–52. 10.1016/S1474-4422(21)00214-3.

[24] Janelidze S, Bali D, Ashton NJ, Barthélemy NR, Vanbrabant J, Stoops E, et al. Head-to-head comparison of 10 plasma phospho-tau assays in prodromal Alzheimer’s disease. Brain 2023;146:1592–601. 10.1093/brain/awac333.

[25] Wang Z, Tan L, Gao P, Ma Y, Fu Y, Sun Y, et al. Associations of the A/T/N profiles in PET, CSF, and plasma biomarkers with Alzheimer’s disease neuropathology at autopsy. Alzheimer’s &amp; Dementia 2023;19:4421–35. 10.1002/alz.13413.

[26] Moscoso A, Grothe MJ, Schöll M, for the Alzheimer’s Disease Neuroimaging Initiative. Reduced [18F]flortaucipir retention in white matter hyperintensities compared to normal-appearing white matter. Eur J Nucl Med Mol Imaging 2021;48:2283–94. 10.1007/s00259-021-05195-5.

[27] Dickson DW, Braak H, Duda JE, Duyckaerts C, Gasser T, Halliday GM, et al. Neuropathological assessment of Parkinson’s disease: refining the diagnostic criteria. The Lancet Neurology 2009;8:1150–7. 10.1016/S1474-4422(09)70238-8.

[28] Liu J, Costantino I, Venugopalan N, Fischetti RF, Hyman BT, Frosch MP, et al. Amyloid structure exhibits polymorphism on multiple length scales in human brain tissue. Sci Rep 2016;6:33079. 10.1038/srep33079.

[29] Araki K, Yagi N, Aoyama K, Choong C-J, Hayakawa H, Fujimura H, et al. Parkinson’s disease is a type of amyloidosis featuring accumulation of amyloid fibrils of α-synuclein. Proc Natl Acad Sci USA 2019;116:17963–9. 10.1073/pnas.1906124116.

[30] Bashit AA, Nepal P, Connors T, Oakley DH, Hyman BT, Yang L, et al. Mapping the Spatial Distribution of Fibrillar Polymorphs in Human Brain Tissue. Front Neurosci 2022;16:909542. 10.3389/fnins.2022.909542.

[31] Suresh K, Dahal E, Badano A. Amyloid models for quantitative x‐ray brain amyloid imaging. Alzheimer’s &amp; Dementia 2023;19:e075380. 10.1002/alz.075380.

[32] Suresh K, Dahal E, Badano A. Synthetic β-sheets mimicking fibrillar and oligomeric structures for evaluation of spectral X-ray scattering technique for biomarker quantification. Cell & Bioscience 2024.

[33] Lake JA. An iterative method of slit-correcting small angle X-ray data. Acta Crystallographica 1967;23:191–4. 10.1107/S0365110X67002440.

[34] Dahal E, Ghammraoui B, Badano A. Characterization of materials embedded in thick objects using spectral small-angle x-ray scattering. J Phys D: Appl Phys 2020;53:245302. 10.1088/1361-6463/ab8248.

[35] Dahal E, Ghammraoui B, Ye M, Smith JC, Badano A. Label-free X-ray estimation of brain amyloid burden. Sci Rep 2020;10:20505. 10.1038/s41598-020-77554-5.

[36] Suresh K, Sharma DK, Chulliyil R, Sarode KD, Kumar VR, Chowdhury A, et al. Single-Particle Tracking To Probe the Local Environment in Ice-Templated Crosslinked Colloidal Assemblies. Langmuir 2018;34:4603–13. 10.1021/acs.langmuir.7b04120.

[37] De Felici M, Felici R, Ferrero C, Tartari A, Gambaccini M, Finet S. Structural characterization of the human cerebral myelin sheath by small angle x-ray scattering. Phys Med Biol 2008;53:5675–88. 10.1088/0031-9155/53/20/007.

[38] O’Brien JS, Sampson EL. Lipid composition of the normal human brain: gray matter, white matter, and myelin. Journal of Lipid Research 1965;6:537–44. 10.1016/S0022-2275(20)39619-X.

[39] Stroud JC, Liu C, Teng PK, Eisenberg D. Toxic fibrillar oligomers of amyloid-β have cross-β structure. Proceedings of the National Academy of Sciences 2012;109:7717–22. 10.1073/pnas.1203193109.

[40] Anagnostou D, Sfakianaki G, Melachroinou K, Soutos M, Constantinides V, Vaikath N, et al. Assessment of Aggregated and Exosome-Associated α-Synuclein in Brain Tissue and Cerebrospinal Fluid Using Specific Immunoassays. Diagnostics 2023;13:2192. 10.3390/diagnostics13132192.

[41] Roberts BR, Lind M, Wagen AZ, Rembach A, Frugier T, Li Q-X, et al. Biochemically-defined pools of amyloid-β in sporadic Alzheimer’s disease: correlation with amyloid PET. Brain 2017;140:1486–98. 10.1093/brain/awx057.

[42] Karran E, Mercken M, Strooper BD. The amyloid cascade hypothesis for Alzheimer’s disease: an appraisal for the development of therapeutics. Nat Rev Drug Discov 2011;10:698–712. 10.1038/nrd3505.

[43] Näslund J. Correlation Between Elevated Levels of Amyloid β-Peptide in the Brain and Cognitive Decline. JAMA 2000;283:1571. 10.1001/jama.283.12.1571.

[44] Cohen SIA, Linse S, Luheshi LM, Hellstrand E, White DA, Rajah L, et al. Proliferation of amyloid-β42 aggregates occurs through a secondary nucleation mechanism. Proc Natl Acad Sci USA 2013;110:9758–63. 10.1073/pnas.1218402110.

[45] Hainfeld JF, Slatkin DN, Focella TM, Smilowitz HM. Gold nanoparticles: a new X-ray contrast agent. BJR 2006;79:248–53. 10.1259/bjr/13169882.

[46] Guerrero-Ferreira R, Kovacik L, Ni D, Stahlberg H. New insights on the structure of alpha-synuclein fibrils using cryo-electron microscopy. Current Opinion in Neurobiology 2020;61:89–95. 10.1016/j.conb.2020.01.014.

[47] Schweighauser M, Shi Y, Tarutani A, Kametani F, Murzin AG, Ghetti B, et al. Structures of α-synuclein filaments from multiple system atrophy. Nature 2020;585:464–9. 10.1038/s41586-020-2317-6.

[48] Martinez-Valbuena I, Visanji NP, Kim A, Lau HHC, So RWL, Alshimemeri S, et al. Alpha-synuclein seeding shows a wide heterogeneity in multiple system atrophy. Transl Neurodegener 2022;11:7. 10.1186/s40035-022-00283-4.

[49] Schmidt MJ, Langen N, Klumpp S, Nasirimanesh F, Shirvanchi P, Ondreka N, et al. A study of the comparative anatomy of the brain of domestic ruminants using magnetic resonance imaging. The Veterinary Journal 2012;191:85–93. 10.1016/j.tvjl.2010.12.026.

[50] Eaton SL, Murdoch F, Rzechorzek NM, Thompson G, Hartley C, Blacklock BT, et al. Modelling Neurological Diseases in Large Animals: Criteria for Model Selection and Clinical Assessment. Cells 2022;11:2641. 10.3390/cells11172641.

[51] Beale AM, Higgins RJ, Work TM, Bailey CS, Smith MO, Shinka T, et al. MPTP-induced Parkinson-like disease in sheep: clinical and pathologic findings. J Environ Pathol Toxicol Oncol 1989;9:417–28.

[52] Baskin DS, Browning JL, Widmayer MA, Zhu Z-Q, Grossman RG. Development of a model for Parkinson’s disease in sheep using unilateral intracarotid injection of MPTP via slow continuous infusion. Life Sciences 1994;54:471–9. 10.1016/0024-3205(94)00406-4.

[53] Nelson PT, Greenberg SG, Saper CB. Neurofibrillary tangles in the cerebral cortex of sheep. Neuroscience Letters 1994;170:187–90. 10.1016/0304-3940(94)90270-4.

[54] Braak H, Braak E, Strothjohann M. Abnormally phosphorylated tau protein related to the formation of neurofibrillary tangles and neuropil threads in the cerebral cortex of sheep and goat. Neuroscience Letters 1994;171:1–4. 10.1016/0304-3940(94)90589-4.

[55] Reid SJ, Mckean NE, Henty K, Portelius E, Blennow K, Rudiger SR, et al. Alzheimer’s disease markers in the aged sheep (Ovis aries). Neurobiology of Aging 2017;58:112–9. 10.1016/j.neurobiolaging.2017.06.020.

[56] Morton AJ. Large-Brained Animal Models of Huntington’s Disease: Sheep. In: Precious SV, Rosser AE, Dunnett SB, editors. Huntington’s Disease, vol. 1780, New York, NY: Springer New York; 2018, p. 221–39. 10.1007/978-1-4939-7825-0_12.

[57] Guo Y-J, Xiong H, Chen K, Zou J-J, Lei P. Brain regions susceptible to alpha-synuclein spreading. Mol Psychiatry 2022;27:758–70. 10.1038/s41380-021-01296-7.

[58] Hampel H, Hardy J, Blennow K, Chen C, Perry G, Kim SH, et al. The Amyloid-β Pathway in Alzheimer’s Disease. Mol Psychiatry 2021;26:5481–503. 10.1038/s41380-021-01249-0.

[59] Spillantini MG, Goedert M. Tau pathology and neurodegeneration. The Lancet Neurology 2013;12:609–22. 10.1016/S1474-4422(13)70090-5.

[60] Doty RL. Olfactory dysfunction in Parkinson disease. Nat Rev Neurol 2012;8:329–39. 10.1038/nrneurol.2012.80.

[61] Chen F, Liu W, Liu P, Wang Z, Zhou Y, Liu X, et al. α-Synuclein aggregation in the olfactory bulb induces olfactory deficits by perturbing granule cells and granular–mitral synaptic transmission. Npj Parkinsons Dis 2021;7:114. 10.1038/s41531-021-00259-7.

[62] Thal DR, Rüb U, Orantes M, Braak H. Phases of Aβ-deposition in the human brain and its relevance for the development of AD. Neurology 2002;58:1791–800. 10.1212/WNL.58.12.1791.

[63] Lowe VJ, Wiste HJ, Senjem ML, Weigand SD, Therneau TM, Boeve BF, et al. Widespread brain tau and its association with ageing, Braak stage and Alzheimer’s dementia. Brain 2018;141:271–87. 10.1093/brain/awx320.

[64] Yushkevich PA, Muñoz López M, Iñiguez De Onzoño Martin MM, Ittyerah R, Lim S, Ravikumar S, et al. Three-dimensional mapping of neurofibrillary tangle burden in the human medial temporal lobe. Brain 2021;144:2784–97. 10.1093/brain/awab262.

[65] Albers J, Pacilé S, Markus MA, Wiart M, Vande Velde G, Tromba G, et al. X-ray-Based 3D Virtual Histology—Adding the Next Dimension to Histological Analysis. Mol Imaging Biol 2018;20:732–41. 10.1007/s11307-018-1246-3.

[66] Badano A, Dahal E, Smith JC, Ghammraoui B. System and method for in vivo estimation of brain amyloid burden using x-rays, https://patentimages.storage.googleapis.com/40/41/b9/6250a7468527d1/WO2023096869A1.pdf

[67] Smith R, Capotosti F, Schain M, Ohlsson T, Vokali E, Molette J, et al. The α-synuclein PET tracer [18F] ACI-12589 distinguishes multiple system atrophy from other neurodegenerative diseases. Nat Commun 2023;14:6750. 10.1038/s41467-023-42305-3.

[68] Harper L, Bouwman F, Burton EJ, Barkhof F, Scheltens P, O’Brien JT, et al. Patterns of atrophy in pathologically confirmed dementias: a voxelwise analysis. J Neurol Neurosurg Psychiatry 2017;88:908–16. 10.1136/jnnp-2016-314978.

[69] Fitzpatrick AWP, Debelouchina GT, Bayro MJ, Clare DK, Caporini MA, Bajaj VS, et al. Atomic structure and hierarchical assembly of a cross-β amyloid fibril. Proc Natl Acad Sci USA 2013;110:5468–73. 10.1073/pnas.1219476110.

[70] Trzeciak P, Herbet M, Dudka J. Common Factors of Alzheimer’s Disease and Rheumatoid Arthritis— Pathomechanism and Treatment. Molecules 2021;26:6038. 10.3390/molecules26196038.

[71] https://www.accessdata.fda.gov/drugsatfda_docs/label/2023/217388s000lbl.pdf

