## Supporting information for "Label-free and reference region-free X-ray cross-β index for quantifying protein aggregates of neurodegenerative diseases"

**Table S1:** The list of literature reported X-ray scattering/microdiffraction studied protein aggregates associated to various neurodegenerative diseases (NDDs). Here we classified the prior studies into two based on the type of X-ray source used (synchrotron/laboratory). We also quantitatively evaluated the scattering profiles of these various reported protein aggregates in terms of area ratio ($a_{sheet}/a_{strand}$), full width half maximum ratio ($\Delta q_{50,sheet}/\Delta q_{50,strand}$) and peak height ratio ($I_{q_c,sheet}/I_{q_c,strand}$) between deconvolved inter-β sheet and inter-β strand peaks. The details of inter-β sheet and inter-β strand peak decompositions from composite X-ray spectrum can be found in the main article. To estimate the inter β-strand or inter-β sheet spacings ($d$) from peak center ($q_c$), $d = \frac{2\pi}{q_c}$.

| **X-ray target [reference]** | **Peak center ($q_c$) $nm^{-1}$** | **Area ratio $a_{sheet}/a_{strand}$** | **FWHM ($\Delta q_{50,i}$) ratio $\Delta q_{50,sheet}/\Delta q_{50,strand}$** | **Peak height ($I_{q_c,i}$) ratio $I_{q_c,sheet}/I_{q_c,strand}$** |
|---|---|---|---|---|
| Synchrotron X-ray source | | | | |
| Lewy body (LB)-mouse senile plaque [1] | 6.10<br>13.47 | 0.36 | 0.99 | 0.52 |
| Parkinson's disease (PD)- Lewy body-region of interest (ROI)-1 Patient 1-76 y/o [1] | 5.7<br>12.1 | 0.096 | 0.29 | 0.32 |
| PD- Lewy body-ROI-2 Patient 2-75 y/o [1] | 6.46<br>13.4 | 0.27 | 0.70 | 0.56 |
| PD- Lewy body-ROI 3 Patient 2-75 y/o [1] | 6.13<br>13.37 | 0.27 | 0.38 | 0.60 |
| Amyloid plaque in human Alzheimer's disease (AD) brain tissue [2] | 13.23<br>13.36<br>13.51 | | | |
| Aβ fibril-atomic coordinates from transmission electron microscopy (TEM) [3] | 6.56<br>13.65 | 0.86 | 1.58 | 0.54 |
| tau fibril-atomic coordinates from TEM [3] | 13.36 | | | |
| Aβ fibrils in AD [3] | 6.33<br>13.685 | 0.921 | 2.06 | 0.45 |

| | | | | |
|---|---|---|---|---|
| tau in AD [3] | 7.5<br>15.56 | 0.90 | 0.86 | 1.16 |
| Tau deposits in human Down syndrome tissue [3] | 13.37 | | | |
| Tau deposits in human frontotemporal lobar degeneration tissue [3] | 13.37 | | | |
| glial cytoplasmic inclusions (GCIs) of multiple system atrophy (MSA)-cerebellum-1 [1] | 5.15<br>12.94 | 0.167 | 0.482 | 0.354 |
| MSA GCI-cerebellum-2 [1] | 5.28<br>13.09 | 0.14 | 0.45 | 0.32 |
| MSA GCI-striatum [1] | 4.98<br>12.93 | 0.091 | 0.36 | 0.26 |
| Iowa mutation: synthetic mutant D23N-Aβ40 fibrils [4] | 6.68<br>13.37 | | | |
| Laboratory X-ray source | | | | |
| Tau filaments from frontal cortex of patients with Down's syndrome [5] | 13.37 | | | |
| In-vitro assembled tau filaments [5] | 13.37 | | | |
| Wet centrifuged pellet of paired helical filaments (PHF) [6] | 6.03<br>13.28 | 0.39 | 1.56 | 0.25 |
| Meridional scans of dry partially oriented PHF [6] | 5.87<br>13.27 | 0.20 | 1.42 | 0.14 |
| Equatorial scans of dry partially oriented PHF [6] | 5.79<br>13.12 | 2.23 | 2.43 | 0.92 |
| Dry pellet of amyloid [6] | 5.85<br>13.31 | 0.50 | 1.20 | 0.42 |

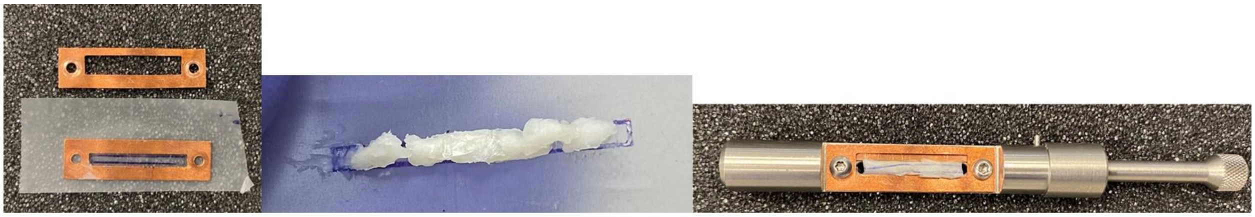

sample holder

oligomer model layered on top of tissue phantom

sample wrapped in adhesive tape and fixed within sample holder slit window

**Figure S1**: Brain tissue phantom preparation using fat with or without the model protein aggregates for wide angle X-ray scattering (WAXS) measurements. The samples are wrapped in adhesive tape and sealed within the rectangular slit shaped sample holder (slit width 2.8 mm and length 20 mm) to ensure the whole sample is within the incident X-ray path and exposed to X-rays.

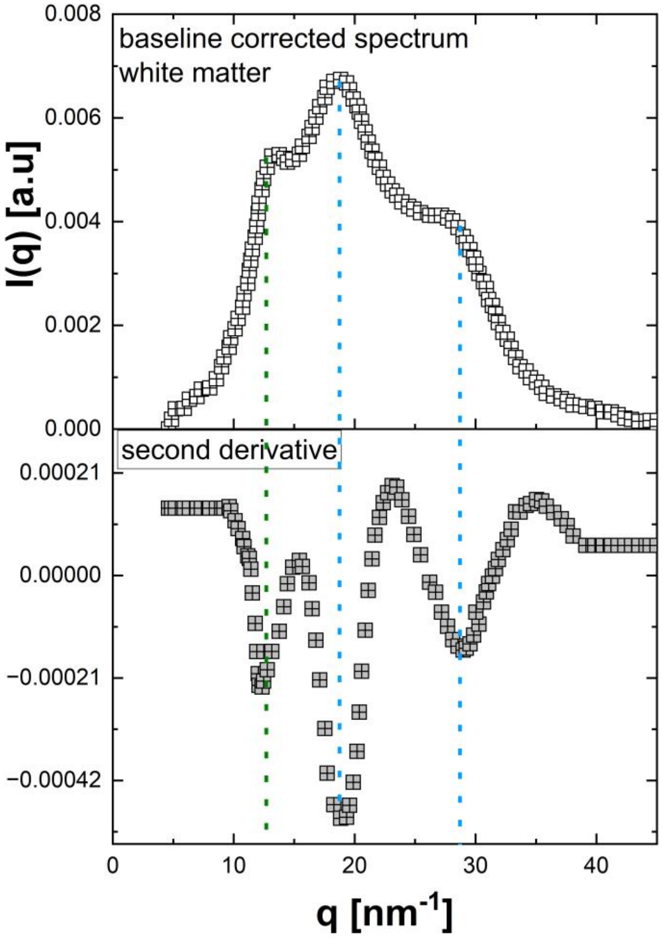


**Figure S2**: Synchrotron X-ray source based WAXS profile of human brain white matter tissues. The data was adopted with the permission from reference [7]. The centers of each peak in the composite scattering spectrum were determined either as positions of local maxima in the spectrum or as minima in the second derivative of the spectrum.

**Table S2:** The weight proportions between tissue phantom material and oligomer model for each blend formulation is reported below.

| **Formulation in terms of $f_{tissue}/f_{agg}$** | **Amount of phantom material (mg)** | **Amount of oligomer model (mg)** |
|---|---|---|
| 1/0 | 28 | 0 |
| 0.997/0.003 | 28 | 0.1 |
| 0.995/0.005 | 28 | 0.15 |
| 0.982/ 0.018 | 28 | 0.5 |
| 0.933/0.067 | 28 | 2 |

| | | |
|---|---|---|
| 0.848/0.152 | 28 | 5 |
| 0.8/0.2 | 28 | 7 |

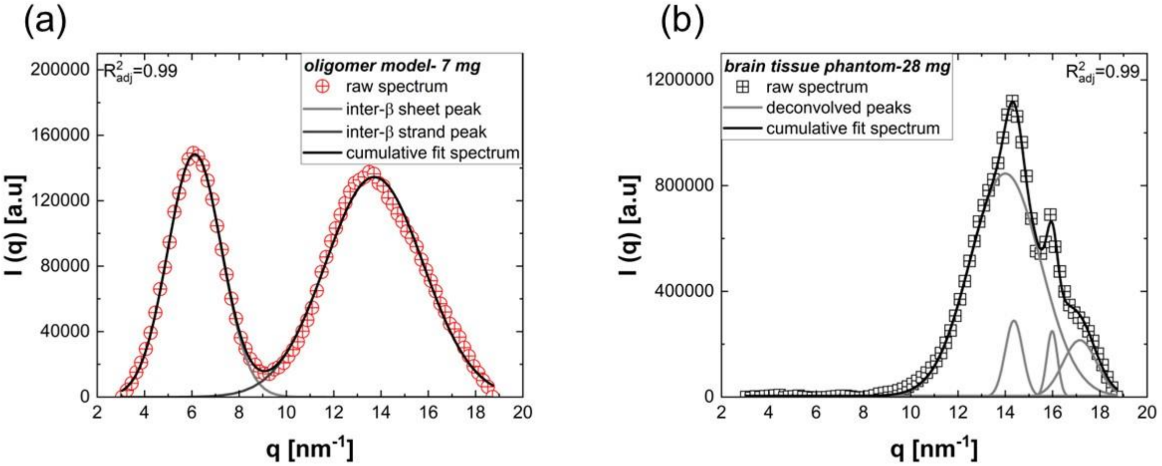


**Figure S3**: The material decomposition of WAXS profiles collected for (a) 7 mg oligomer and (b) 28 mg brain tissue phantom. The goodness of fit for each spectrum is reported in terms of $R^2_{adj}$.

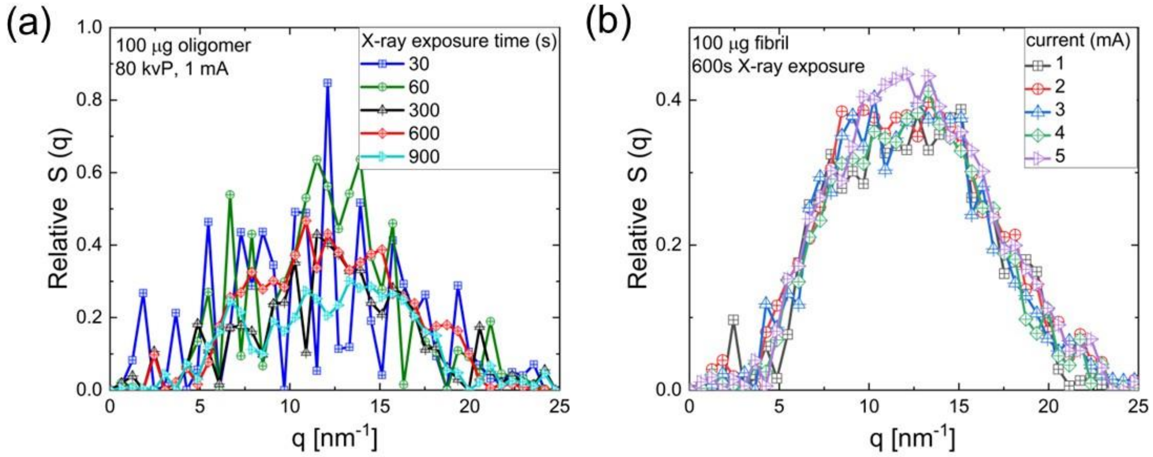


**Figure S4**: The effect of (a) X-ray exposure time on the scattering profile of 100 µg oligomer for a fixed voltage (80 kVp) and current (1 mA) and (b) current on the scattering profile for fixed voltage (80 kVp) and X-ray exposure time (600 s).

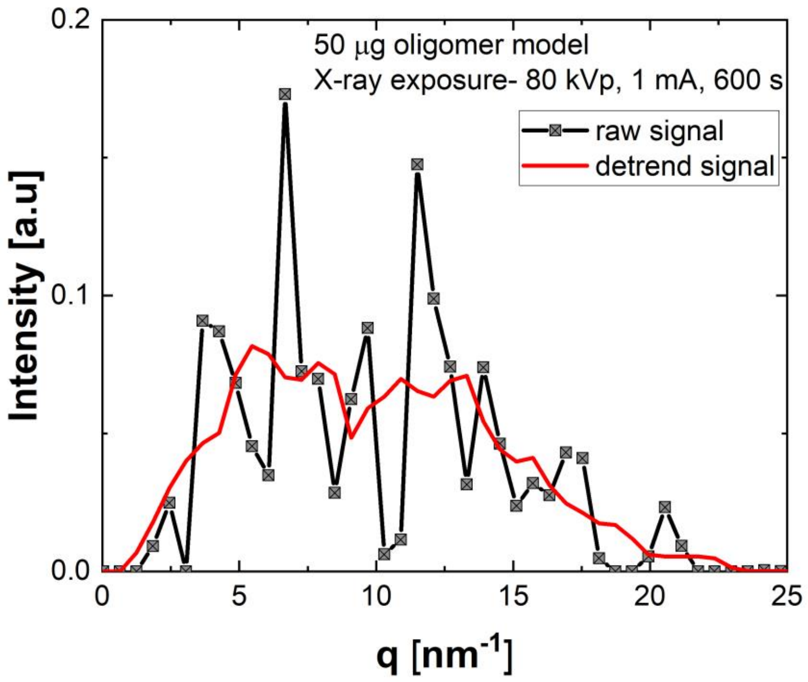


**Figure S5:** The raw X-ray signal of 50 μg model oligomer collected at 80 kVp, 1 mA for 600 s. The raw signal was wavelet denoised using Daubechies wavelet as discussed in section 2.4 of the main article to separate the trend and detrend signals and to estimate the signal to noise ratio.

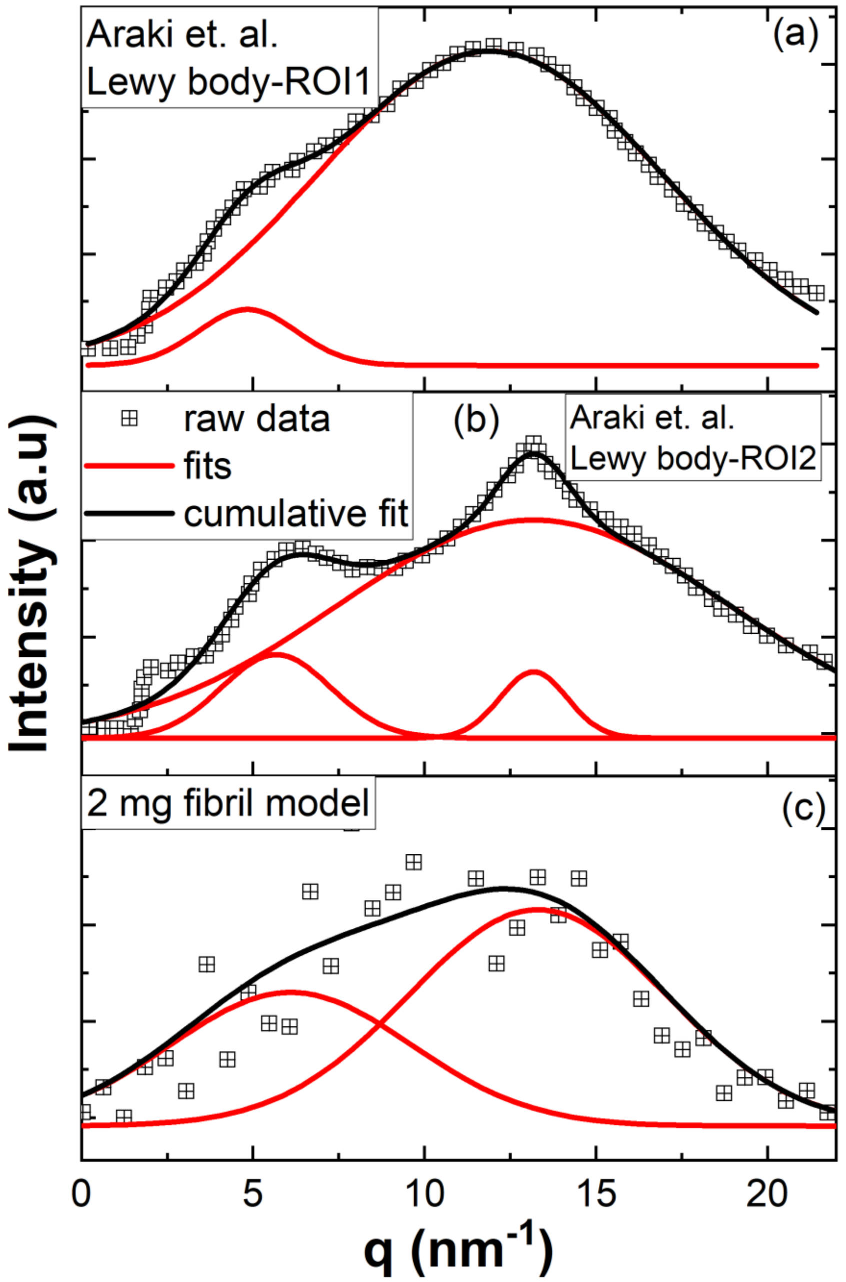


**Figure S6**: X-ray scattering profiles of Lewy bodies formed from α-synuclein aggregates in Parkinson's disease (a) patient 1 (76-year-old man) and (b) patient 2 (75-year-old man). The data (a) and (b) are adopted with the permission from reference [1] (c) WAXS profile of 2 mg LGAB fibrillar model. The raw experimental data are shown with the symbols, red line curves show the decomposed inter β-sheet and inter-β strand peaks and black line curve is the cumulative fit for each composite spectrum.

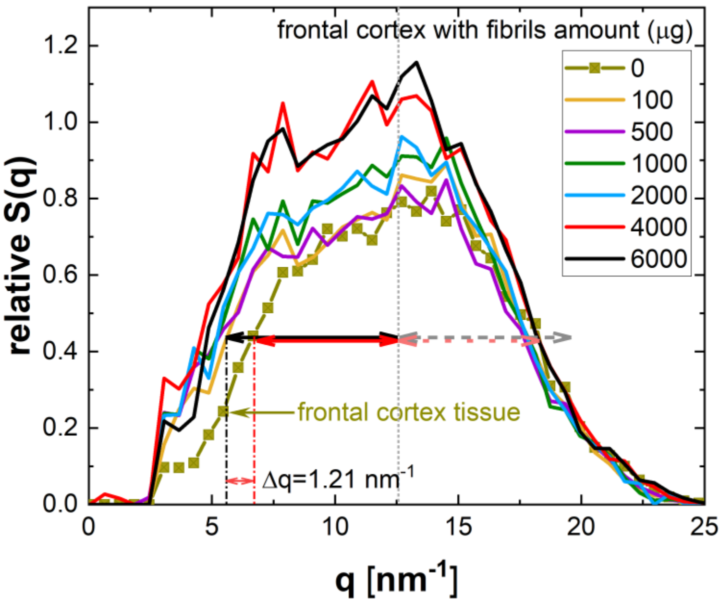


**Figure S7:** The WAXS spectra of sheep brain frontal cortex tissue with varying fibril amount from 100 µg to 6 mg. The increase in spectrum width is shown as the difference in the $q$ position for a specific composite spectrum amplitude with the addition of 100 µg fibril to the healthy frontal cortex tissue.

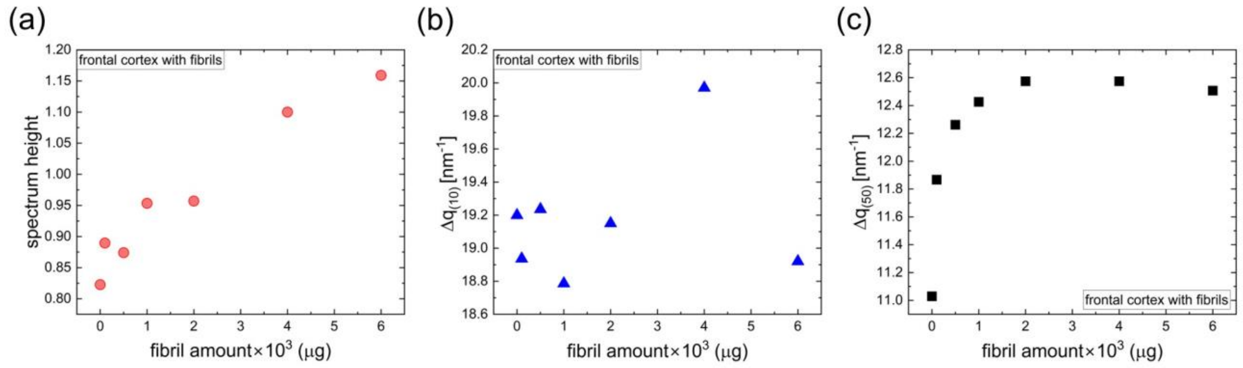


**Figure S8:** The composite spectrum properties of frontal cortex with varying fibrillar amount. (a) Variation of composite spectrum (a) maximum amplitude, (b) full width tenth maximum (FWTM, $\Delta q_{(10)}$) and (c) full width half maximum (FWHM, $\Delta q_{(50)}$) as a function of fibril amount in the region of interest (ROI) of a tissue.

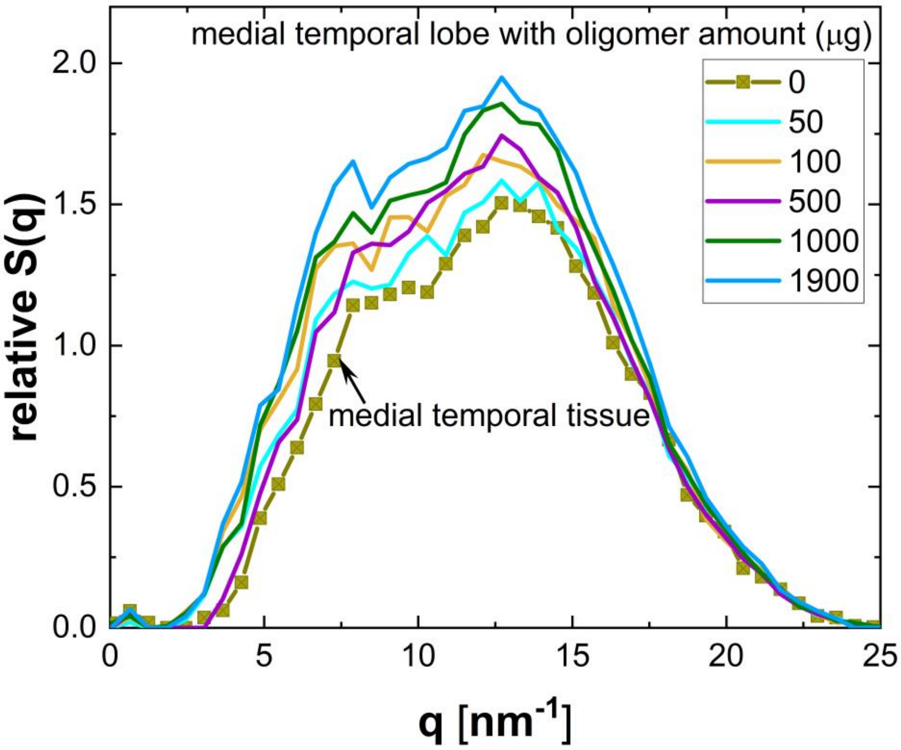


**Figure S9**: The WAXS profiles of the tissue from medial temporal lobe before and after the addition of varying quantities of oligomer.

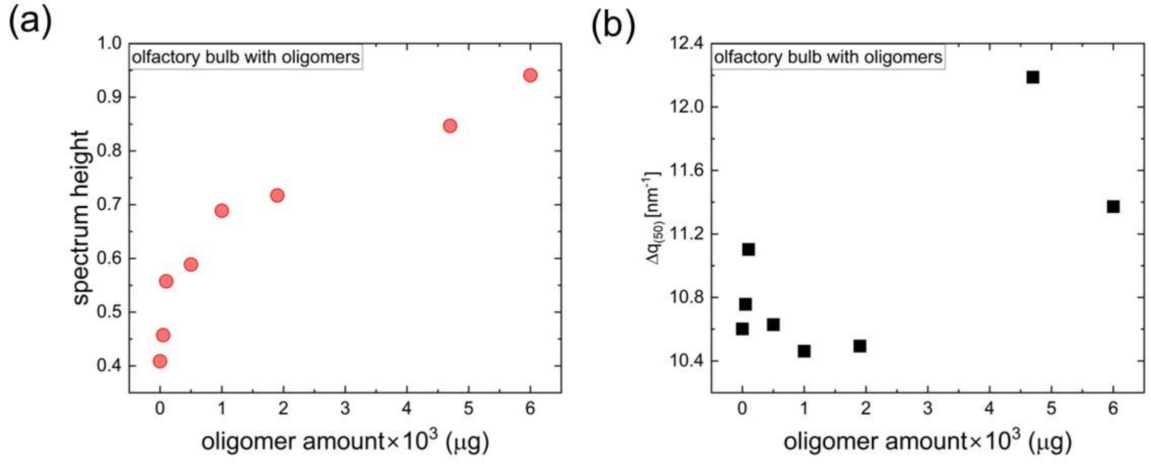


**Figure S10**: The characteristics of composite spectrum of olfactory bulb before and after the addition of varying quantities of oligomer to the same region of interest (ROI) of the tissue. The composite spectrum (a) maximum amplitude and (b) full width half maximum (FWHM, $\Delta q_{(50)}$) as a function of oligomer amount.

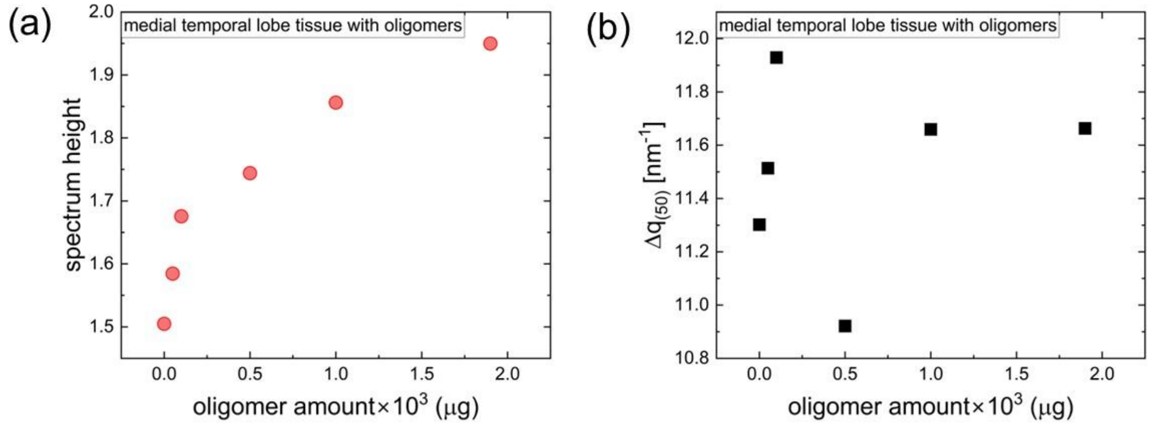


**Figure S11**: The characteristics of composite spectrum of tissue from medial temporal lobe region before and after the addition of varying quantities of oligomer to the same region of interest (ROI) of the tissue. The composite spectrum (a) maximum amplitude and (b) full width half maximum (FWHM, $\Delta q_{(50)}$) as a function of oligomer amount.

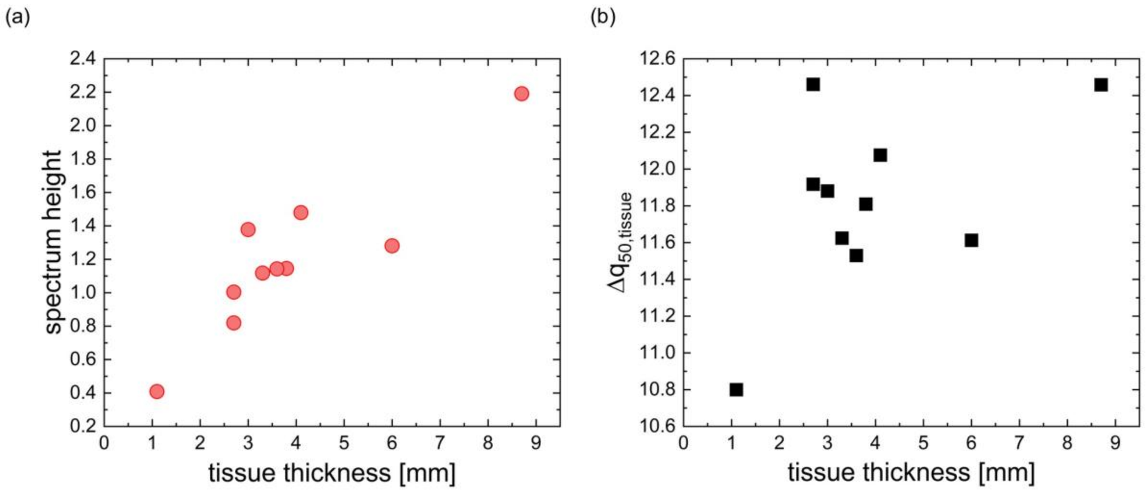


**Figure S12**: The WAXS spectrum characteristics of different tissue regions which are marked in Figure 6a in the main article. The variation of composite spectrum (a) maximum amplitude and (b) full width half maximum (FWHM, $\Delta q_{(50)}$) as a function of tissue thickness. The tissue thickness represents the average of three measurements obtained using a vernier caliper within each tissue ROI.

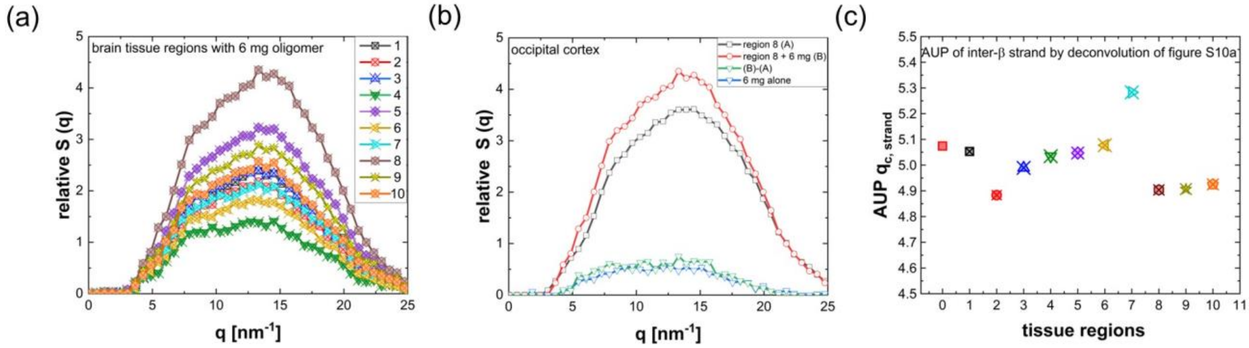


**Figure S13**: (a) The composite WAXS profiles of ten different tissue regions of interest with 6 mg oligomer burden. The tissue regions are marked in Figure 6a in the main article. The WAXS profiles of ROI matched tissues prior to the oligomer addition can be seen as Figure 6b. (b) The WAXS spectrum of the occipital cortex tissue is represented by the black line with symbols, while the occipital cortex tissue with 6 mg oligomer is depicted by the red line with symbols. The green line with symbols illustrates the result of subtracting the red line curve from the black line curve. Additionally, the standard of reference, which is the WAXS profile of 6 mg oligomer alone, is shown by the blue line with symbols. The subtracted profile and standard of reference exhibit comparability. (c) The composite spectra in Figure S13a were decomposed to tissue signal, inter β-sheet and inter β-strand signals of 6 mg oligomer. The AUP of the decomposed inter-β strand signal for oligomer in ten different composite spectra is shown here. The AUP of zeroth tissue region is the value of reference standard.

### REFERENCES


[1] Araki K, Yagi N, Aoyama K, Choong C-J, Hayakawa H, Fujimura H, et al. Parkinson's disease is a type of amyloidosis featuring accumulation of amyloid fibrils of α-synuclein. Proc Natl Acad Sci USA 2019;116:17963–9. https://doi.org/10.1073/pnas.1906124116.

[2] Liu J, Costantino I, Venugopalan N, Fischetti RF, Hyman BT, Frosch MP, et al. Amyloid structure exhibits polymorphism on multiple length scales in human brain tissue. Sci Rep 2016;6:33079. https://doi.org/10.1038/srep33079.

[3] Bashit AA, Nepal P, Connors T, Oakley DH, Hyman BT, Yang L, et al. Mapping the Spatial Distribution of Fibrillar Polymorphs in Human Brain Tissue. Front Neurosci 2022;16:909542. https://doi.org/10.3389/fnins.2022.909542.

[4] Tycko R, Sciarretta KL, Orgel JPRO, Meredith SC. Evidence for Novel β-Sheet Structures in Iowa Mutant β-Amyloid Fibrils. Biochemistry 2009;48:6072–84. https://doi.org/10.1021/bi9002666.

[5] Berriman J, Serpell LC, Oberg KA, Fink AL, Goedert M, Crowther RA. Tau filaments from human brain and from in vitro assembly of recombinant protein show cross-β structure. Proc Natl Acad Sci USA 2003;100:9034–8. https://doi.org/10.1073/pnas.1530287100.

[6] Kirschner DA, Abraham C, Selkoe DJ. X-ray diffraction from intraneuronal paired helical filaments and extraneuronal amyloid fibers in Alzheimer disease indicates cross-beta conformation. Proc Natl Acad Sci USA 1986;83:2776–2776. https://doi.org/10.1073/pnas.83.8.2776-b.

[7] De Felici M, Felici R, Ferrero C, Tartari A, Gambaccini M, Finet S. Structural characterization of the human cerebral myelin sheath by small angle x-ray scattering. Phys Med Biol 2008;53:5675–88. https://doi.org/10.1088/0031-9155/53/20/007.
